# Temporal pattern recognition in the human brain: a dual simultaneous processing

**DOI:** 10.1101/2021.10.21.465263

**Authors:** L. Bonetti, E. Brattico, S.E.P. Bruzzone, G. Donati, G. Deco, D. Pantazis, P. Vuust, M.L. Kringelbach

## Abstract

Pattern recognition is a major scientific topic. Strikingly, while machine learning algorithms are constantly refined, the human brain emerges as an ancestral biological example of such complex procedure. However, how it transforms sequences of single objects into meaningful temporal patterns remains elusive. Using magnetoencephalography (MEG) and magnetic resonance imaging (MRI), we discovered and mathematically modelled an inedited dual simultaneous processing responsible for pattern recognition in the brain. Indeed, while the objects of the temporal pattern were independently elaborated by a local, rapid brain processing, their combination into a meaningful superordinate pattern depended on a concurrent global, slower processing involving a widespread network of sequentially active brain areas. Expanding the established knowledge of neural information flow from low- to high-order brain areas, we revealed a novel brain mechanism based on simultaneous activity in different frequency bands within the same brain regions, highlighting its crucial role underlying complex cognitive functions.

## Introduction

Pattern recognition has gathered a large interest across all scientific fields. Indeed, as a consequence of technological developments, nowadays scientists can rely on constantly growing amount of data and computational power ^1,2^. This has introduced new exciting opportunities, pushing research to seek complex patterns that emerged in a variety of different fields, spanning from quantum physics ^3^ to weather forecast ^4^, animal and human behavior ^5,6^, and medical imaging ^7,8^.

Strikingly, while computer science constantly refines machine learning algorithms and artificial intelligence for pattern recognition, neuroscience proposes the human brain as an ancestral biological example of such complex procedure ^9–11^. Indeed, to guarantee survival the brain urges to invariably learn and recognize patterns. Among others, brain research findings on synchronous visual patterns detection provided major advances about the brain mechanisms underlying face and object recognition ^12–15^. These studies showed the key role of fusiform gyrus for face recognition and highlighted the cascade of events from primary visual cortex to higher-order associative areas underlying processing and recognition of visual objects ^16,17^.

When investigating the neural responses to objects arranged over time, it has been demonstrated that the brain is able to automatically detect regularities in temporal patterns (sequences) of objects, even at a pre-attentive level ^18–20^. This research, largely carried out in the auditory domain ^18^, discovered automatic event-related potentials/fields (ERP/F) to deviant and standard stimuli such as N100, mismatch negativity (MMN) ^18,19^ and early right-anterior negativity (ERAN) ^21,22^.

Additional studies in the context of auditory neuroscience and memory for sounds arranged over time highlighted a network of brain areas supposedly involved in storing and retrieving acoustic information, comprising auditory cortex, inferior frontal gyrus and hippocampus ^23–26^. These works, which mainly employed functional magnetic resonance imaging (fMRI), returned pivotal results on auditory memory, and increased our knowledge on how the brain actively manipulates sounds and auditory information extended over time. Nonetheless, they did not unravel the fast-scale brain spatial-temporal dynamics responsible for conscious temporal pattern encoding and recognition.

Thus, despite decades of advancements in neuroscience, several open questions remain. In particular, how does the brain transform sequences of single objects (local processing) in meaningful temporal patterns (global processing) which are accessible to human awareness? What are the core spatial-temporal brain mechanisms behind temporal pattern recognition?

To address such crucial questions, in our study we investigated the brain activity during conscious recognition of auditory patterns extended over time.

## Results

### Experimental design and MEG sensors analysis

To elaborate the ideal experimental design and stimuli, we employed the human activity that mostly acquires meaning by unfolding over time, namely music ^27^. Indeed, after requesting 70 participants to memorize a full musical piece composed by J.S. Bach, we presented them with a set of melodic excerpts taken from the piece and a series of new variations thereof (Fig. 1A). Those excerpts represented temporal patterns built by five objects (musical tones) that were listened by the participants and labelled as ‘previously memorized’ (M) or ‘novel’ (N), using a response pad. The experiment took place while their brain activity was measured through magnetoencephalography (MEG), a powerful machine which records neural activity with excellent time resolution (1-ms precision). In the first place, after preprocessing the MEG data (see Fig. 1B and Methods for details), we analyzed the brain activity underlying recognition of M vs N by using univariate tests for each MEG sensor and time-point and corrected for multiple comparisons with cluster-based Monte-Carlo simulations (MCS). This procedure returned a large significant cluster (*p* < .001, cluster size k = 2117, mean *t-value* = 3.29, time = 0.547 – 1.180 s), showing stronger brain activity for M vs N. Moreover, the brain activity recorded over the MEG channels forming such significant cluster outlined a timeseries which presented two main frequency components. As shown in Fig. S1A, the faster frequency component peaked after the presentation of each of the objects forming the pattern, while the slower frequency component accompanied the whole pattern, peaking at its end. This evidence was further supported by the computation of complex Morlet wavelet transform on all MEG sensor data, which highlighted the main contribution of 1Hz and 4Hz to the MEG signal recorded during the task (Fig. S1B). Thus, our following analyses focused on two frequency bands defined around those main frequencies: 0.1-1Hz and 2-8Hz. These bands roughly corresponded to the well-known brain waves called delta and theta, respectively ^28^, and were also coherent with results reported by Bonetti and colleagues ^29,30^. Importantly, we hypothesized that they indexed the two main processes involved in our experimental task: processing of single objects forming the temporal pattern (*i - local processing*) and recognizing the temporal pattern as a comprehensive superordinate object (*ii - global processing*).

**Fig. 1.**
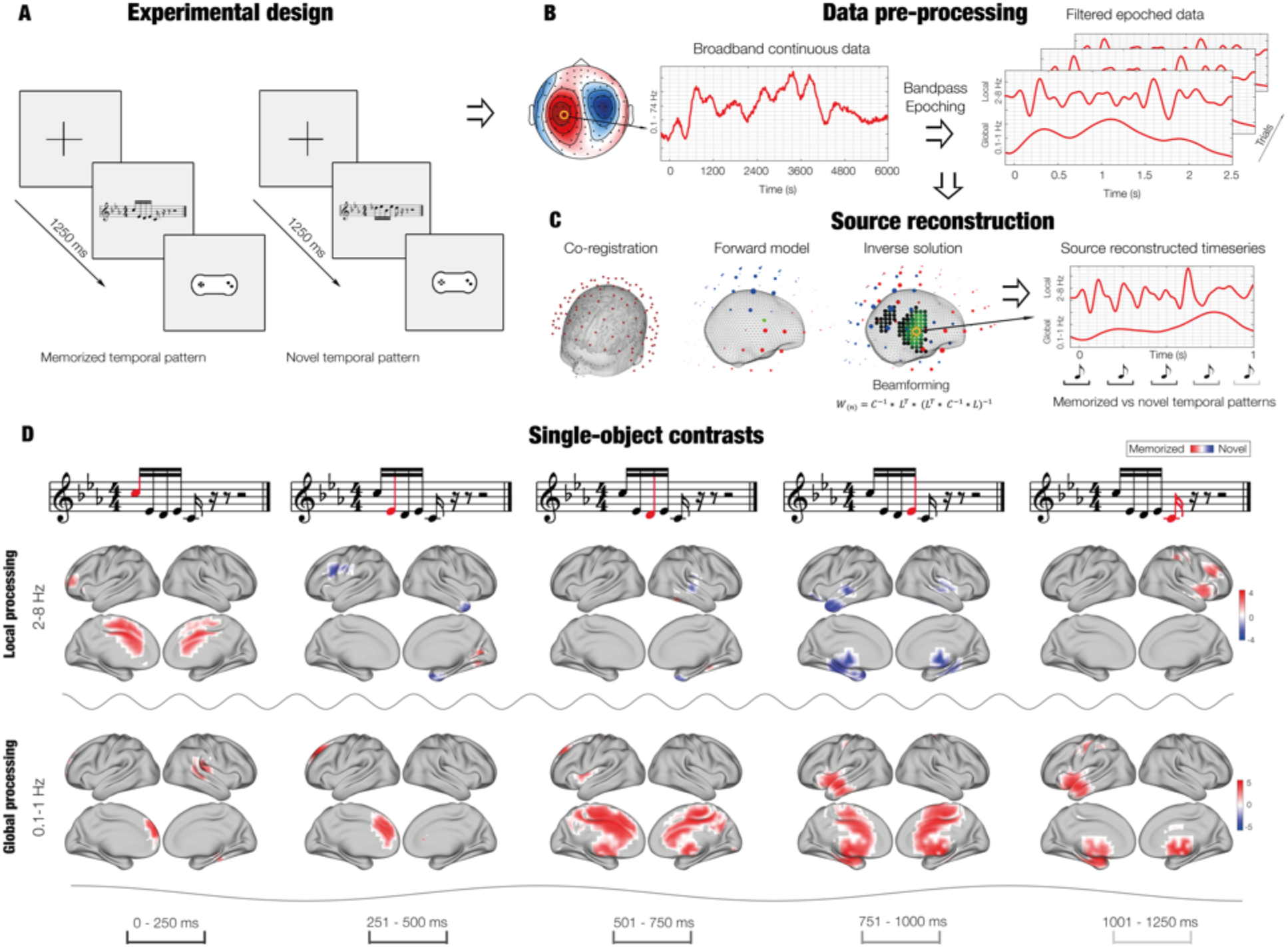
Experimental design, source reconstruction and single-object contrasts. (**A**) After listening to a full musical piece composed by J.S. Bach, participants were presented with a set of melodic excerpts taken from the piece and a series of new variations thereof. Those excerpts represented temporal patterns built by five objects (musical tones) that where labelled by the participants as ‘previously memorized’ (M) or ‘novel’ (N) using a response pad. (**B**) During the task, participant’s brain activity was recorded through MEG. The neural data was preprocessed, bandpass-filtered in two frequency bands (0.1-1Hz and 2-8Hz) and epoched. (**C**) Graphical depiction of source reconstruction, computed independently for the two frequency bands considered in the study. Notably, the slower band (0.1-1Hz) indexed the recognition of the whole pattern (global processing), while the faster band (2-8Hz) showed the neural responses to each object of the pattern (local processing). (**F**). Contrasts revealed stronger brain activity for M vs N in 0.1-1Hz (red), especially for third, fourth and fifth objects. Such difference was localized in a large brain network comprising cingulum, inferior temporal cortex, frontal operculum, insula and hippocampal areas. Conversely, contrasts for 2-8Hz returned an overall stronger activity for N vs M (blue), especially in the auditory cortex.

### Source reconstructed brain activity and single-object analysis

MEG is a powerful tool to record whole-brain activity with excellent temporal resolution. However, the investigation of neural activity also requires reliable spatial parameters. To achieve such goal, we performed the widely adopted procedure named source reconstruction, implemented through a beamforming algorithm. As shown in Fig. 1B, we band-pass filtered the MEG continuous data in the previously mentioned frequency bands (0.1-1Hz, delta, and 2-8Hz, theta). Then, independently for the two bands, the MEG data was epoched and co-registered (Fig. 1C) with the structural images of the participants’ brains (structural weighted T1) obtained from magnetic resonance imaging (MRI). Finally, using beamforming, we reconstructed the neural sources of the MEG signal in an 8-mm space, returning 3559 brain sources (voxels) and the timeseries showing their activity over time (see Fig. 1C and Methods for details).

The reconstructed brain activity of each participant was submitted to first-level analysis, which was conducted employing general linear models (GLMs). Such models were computed for each time-point and brain voxel, returning the main effect (contrasts of parameters estimate (COPEs)) of M and N as well as their contrast ^31^. These results were submitted to second level (group-level) analysis, employing one-sample t-tests with spatially smoothed variance obtained with a Gaussian kernel (full-width at half-maximum: 50 mm). This analysis returned the group-level statistics over all participants for each time-point and brain voxel, independently for our two frequency bands.

Then, we aimed to investigate the brain activity underlying the local processing of each of the objects forming the temporal pattern (theta) as well as the simultaneous global processing of the whole pattern (delta). Thus, we computed 10 (five tones x two frequency bands) cluster-based MCS on the group-level results averaged over the five time-windows corresponding to the duration of the musical tones. The MCS analysis comprised 1000 permutations and a cluster forming threshold of *p* < .05 (from the group-level analysis). Since we computed this analysis 10 times, we corrected for multiple comparisons by dividing the standard MCS *α* level (= .05) by our 10 comparisons, resulting in an updated MCS *α* = .005 (i.e. clusters of significant group-level results in the original data were significant if their sizes were larger than the 99.5% of the cluster sizes of the permuted data; see Methods for additional details).

Notably, different results emerged for the two frequency bands. Brain activity for delta was stronger for M vs N, especially during processing of the last three objects of the pattern. As depicted in Fig. 1D and S2, such activity delineated a widespread brain network underlying the global processing of the pattern, involving brain regions related to memory and evaluative processes such as cingulate gyrus, hippocampus, insula, frontal operculum and inferior temporal cortex (MCS *p* < .001). Conversely, brain activity for theta was overall stronger for N vs M and mainly involved auditory cortices (MCS *p* < .001). Statistics of the peak significant brain voxels for both frequencies are reported in Table 1, while extensive results are described in **Table S1**.

**Table 1.** Peak brain activity underlying recognition of each object (musical tone) of the temporal sequences. Brain areas refer to the automatic anatomic labelling (AAL) parcellation labels, while t indicates the t-value obtained by contrasting known vs unknown temporal sequences.

| 0.1 – 1 Hz |  |  |  |  |  | 2 – 8 Hz |  |  |  |  |  |
| --- | --- | --- | --- | --- | --- | --- | --- | --- | --- | --- | --- |
| Brain area | Hemisph | t | MNI coordinates |  |  | Brain area | Hemisph | t | MNI coordinates |  |  |
|  |  |  | x | y | z |  |  |  | x | y | z |
| **Tone 1** |  |  |  |  |  |  |  |  |  |  |  |
| Rolandic Ope | R | 4.44 | 42 | -30 | 16 | Cing Mid | R | 4.48 | 2 | 2 | 40 |
| Heschl | R | 4.26 | 42 | -30 | 8 | Cing Mid | R | 4.23 | 2 | 10 | 40 |
| Temporal Sup | R | 4.16 | 50 | -30 | 16 | Cing Mid | L | 4.17 | -6 | 2 | 40 |
| Temporal Sup | R | 4.04 | 42 | -38 | 16 | Cing Mid | R | 4.16 | 2 | 10 | 32 |
| **Tone 2** |  |  |  |  |  |  |  |  |  |  |  |
| Frontal Sup | L | 4.00 | -14 | 34 | 40 | Tempr Pol Sup | R | -3.88 | 34 | 10 | -32 |
| Frontal Sup | L | 3.98 | -14 | 34 | 32 | Tempr Pol Sup | R | -3.46 | 26 | 10 | -32 |
| Frontal Sup | L | 3.92 | -14 | 26 | 40 | Front Inf Ope | L | -3.45 | -38 | 2 | 24 |
| Frontal Sup | L | 3.78 | -14 | 42 | 40 | Tempr Pol Mid | R | -3.37 | 42 | 10 | -32 |
| **Tone 3** |  |  |  |  |  |  |  |  |  |  |  |
| Precuneus | R | 3.89 | 2 | -46 | 48 | Temporal Sup | R | -3.30 | 50 | -22 | 8 |
| Cing Mid | R | 3.80 | 2 | -38 | 48 | Temporal Sup | R | -3.19 | 58 | -22 | 8 |
| Cing Mid | R | 3.62 | 2 | -22 | 48 | Temporal Sup | R | -2.78 | 50 | -22 | 0 |
| Cing Mid | R | 3.60 | 2 | -30 | 48 | Heschl | R | -2.67 | 42 | -22 | 8 |
| **Tone 4** |  |  |  |  |  |  |  |  |  |  |  |
| Temporal Mid | L | 5.05 | -46 | -6 | -16 | ParaHippocamp | L | -3.89 | -22 | -30 | -16 |
| Insula | L | 4.93 | -38 | -6 | -8 | ParaHippocamp | L | -3.86 | -30 | -30 | -16 |
| Temporal Mid | L | 4.81 | -46 | -14 | -16 | Tempr Pol Mid | L | -3.74 | -46 | 10 | -32 |
| Cing Mid | R | 4.76 | 2 | -6 | 40 | Tempr Pol Mid | L | -3.70 | -38 | 10 | -32 |
| **Tone 5** |  |  |  |  |  |  |  |  |  |  |  |
| Insula | L | 5.48 | -38 | 2 | -8 | Front Inf Tri | R | 3.41 | 42 | 26 | 24 |
| Putamen | L | 5.27 | -30 | 2 | -8 | Putamen | R | 3.36 | 34 | 2 | 0 |
| Temporal Mid | L | 5.26 | -46 | -6 | -16 | Insula | R | 3.28 | 42 | 10 | 0 |
| Temp Pol Mid | L | 5.24 | -46 | 2 | -16 | Postcentral Gyr | R | 3.23 | 34 | -30 | 48 |

### K-means functional clustering

Although returning large significant brain networks and valuable information, contrasting the brain activity in response to each object of the pattern did not fully benefit from the excellent temporal resolution of the MEG data and underestimated brain processes happening in between two or more objects of the pattern. Furthermore, computing contrasts for each time-point and brain voxel returns a large amount of data which is partially redundant and sometimes not straightforward to understand and ideal to mathematically model. Indeed, several brain sources are highly correlated because of both biological reasons involving large populations of neurons generating the signal and artificial source leakage introduced during the source reconstruction^32^. Thus, defining a functionally based parcellation of the brain is of great importance when aiming to synthesize and mathematically describe the spatial extent of the active brain sources as well as their activity over time.

We used a so-called k-means functional clustering, which relies on the combination of k-means clustering computed both on functional and spatial information of each of the reconstructed brain voxel timeseries (Fig. 2A). In brief, first this procedure clusters the 3559 brain voxels in *n* functional parcels according to basic functional features, such as the absolute value of the peaks of the voxels timeseries (Fig. 2B, right) or their corresponding time-indices (Fig. 2B, left). Notably, in our study, delta presented different peaks of activity shifted over time and thus was clustered considering the time-indices of such peaks (Fig. 2B, left). Differently, theta showed very correlated activity and was therefore clustered using the absolute values of such peak activity (Fig. 2B, right). Second, each of the returned *n* functional parcels is further divided according to the spatial information (three-dimensional coordinates) of the brain voxels that belong to it (Fig. 2C). The whole procedure provides a final parcellation and the corresponding timeseries based on both the functional and spatial profile of each of the reconstructed 3559 brain voxels (examples of this procedure are depicted in Fig. S3, S4 and S5 and described in detail in the Methods and in **Tables S2, S3, S4** and **S5, S6**).

**Fig. 2.**
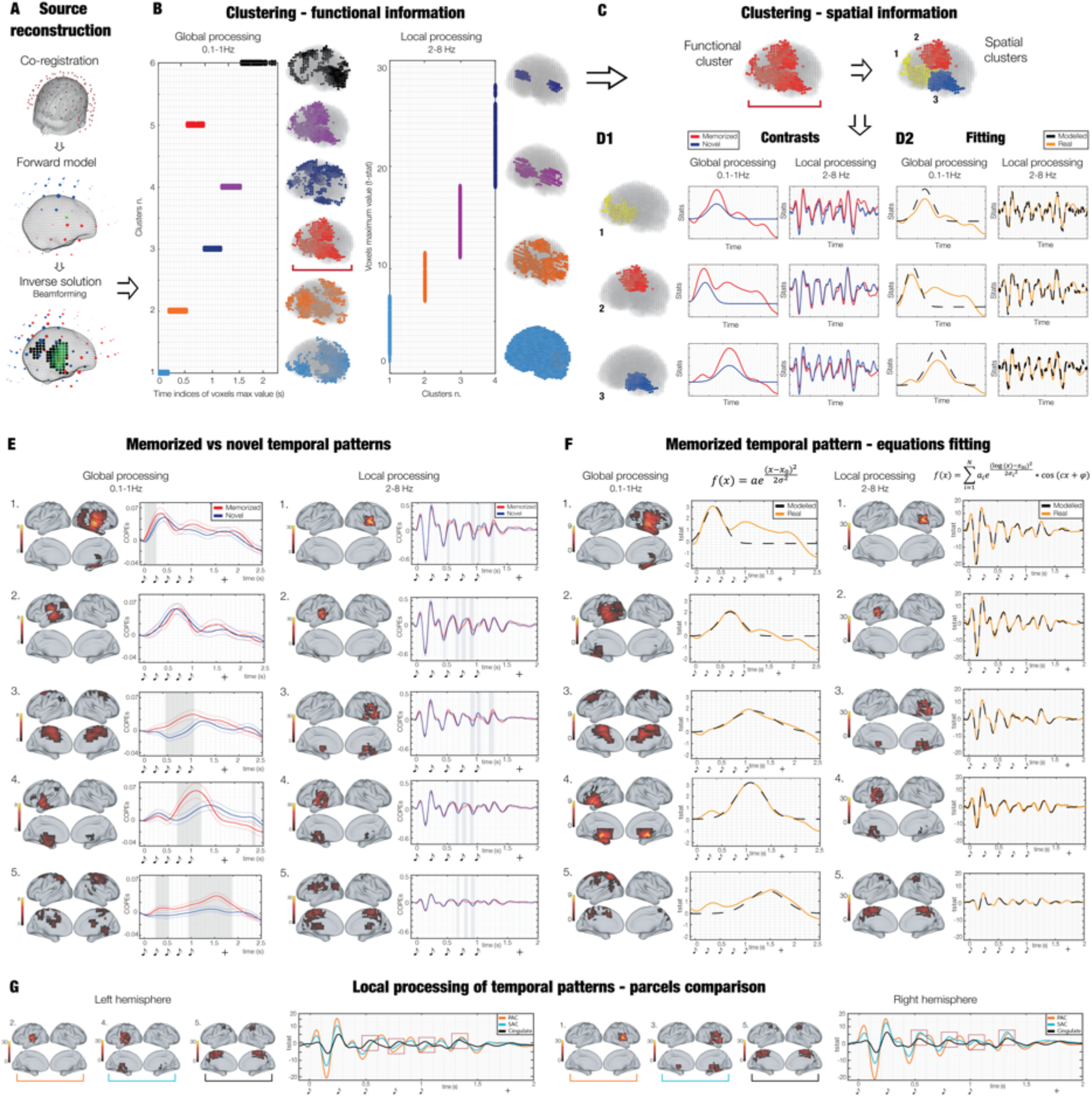
Source reconstruction, k-means functional clustering, contrasts and mathematical modelling. (**A**), (**B**), (**C**) provide a graphical depiction of the methods used, while (**E**), (**F**), (**G**) show actual results. (**A**) Graphical depiction of the source reconstruction. (**B**) A functional parcellation of the brain based on the activity recorded during the task was estimated. First, k-means clustering was computed on functional information of each brain voxel timeseries. Regarding the slower band (0.1-1Hz, indexing the global processing of the pattern), clustering was computed on the time indices of the maximum values of the voxels. Conversely, for the faster band (2-8Hz, indexing the local processing of each object of the pattern), clustering was performed on the maximum values of the voxels. This procedure returns a set of functional parcels. (**C**) A second series of k-means clustering was computed on the spatial properties of each of the functional parcels described in (**B**). Here, for illustrative purposes, we show only one functional parcel (outlined by the red bracket). Such procedure returned a set of new final parcels with the corresponding timeseries taking into account both functional and spatial information of each of the brain voxels. (**D1**) Contrasts between memorized (M) and novel (N) temporal patterns were computed for each parcel and frequency band. (**D2**) Gaussian and sinusoidal functions were fitted to the timeseries of the parcels computed for M only, using the non-linear least square algorithm. (**E**) Contrasts between M vs N temporal patterns for the main functional brain parcels. (**F**) Real and predicted timeseries for M computed by fitting the mathematical equation depicted in the top column to the parcels timeseries. (**G**) Deepening on three main parcels (primary auditory cortex (PAC), secondary auditory cortex and hippocampal areas (SAC) and cingulate) of the local processing. The image highlights the different behavior of PAC vs SAC and Cingulate, especially in the right hemisphere. It shows that higher-order areas (SAC and Cingulate) are more implicated than lower order ones (PAC) in the generation of the P300 component in response to each sound of the pattern (as outlined by the red squares). In (**E**), (**F**), (**G**) graphical depiction of musical tones indicates the onset of the objects forming the temporal pattern, while the ‘+’ shows the mean reaction time of participants’ response. Colorbars refer to the t-values obtained from second-level analyses.

Here, we wanted to contrast the brain activity of M vs N over the functionally defined parcels, aiming to integrate our previous statistical analysis. Thus, the k-means functional clustering was performed on the group-level main effects of M and N averaged together. Then, to obtain the main effect of M and N for each parcel and participant, we averaged the first-level main effect of M and N (from the GLMs) over the brain voxels belonging to each of the functional parcels. This resulted in a new timeseries for each participant, functional parcel, and experimental condition (M and N). Such timeseries were submitted to univariate contrasts (M vs N), performed for each parcel and time-point (Fig. 2D1). Once again, the significant results were corrected for multiple comparisons using cluster based MCS (see Methods for details). These analyses were computed independently for the two frequency bands, returning a different picture for global and local temporal pattern brain processing. Similar to our previous analysis, the strongest brain activity in the delta band was detected for M. Remarkably, the current procedure highlighted a series of sequentially active brain parcels accompanying the processing of the temporal pattern, expanding our first analysis. As shown in Figure 2E, the brain presented an initial activity in the right auditory cortex characterized by a slightly stronger power for M vs N (Fig. 2A1, parcel 1: *p* < .001, cluster size k = 39; mean *t-value* = 2.72; time from first object onset: 0-0.25s). Next, we observed neural activity in the left auditory cortex but no significant differences between experimental conditions (Fig. 2E, parcel 2). Starting between the second and third objects and peaking during the fifth object of the temporal pattern, we observed a burst of activity in the cingulate gyrus, which was stronger for M vs N (Fig. 2E, parcel 3: *p* < .001, k = 92; *t-val* = 2.73; time: 0.45-1.05s). With a slight delay, a similar profile emerged for a larger brain parcel comprising insula, the anterior part of the inferior temporal cortex, hippocampus and frontal operculum. Once again, M was largely stronger than N (Fig. 2E, parcel 4: *p* < .001, k = 77; *t-val* = 2.79; time: 0.69-1.19s). Finally, peaking just before the mean reaction time for participants’ categorization of the pattern, a stronger activity in post-central gyrus and sensorimotor cortex was observed for M vs N (Fig. 2E, parcel 5, main cluster: *p* < .001, k = 142; *t-val* = 2.68; time: 0.94-1.88s).

Conversely, the analysis for theta band showed a number of significant clusters of stronger activity for N vs M around the sharp peaks of the timeseries. Notably, compared to our first analysis for the five objects of the temporal pattern, this second procedure clearly outlined the temporal extent of such difference, which corresponded to the last three tones of the temporal sequences. Specifically, such differences involved right (Fig. 2E, parcel 1, main cluster I: *p* < .001, k = 11, *t-val* = 3.51; time: 0.89-0.95s; II: *p* < .001, k = 11; *t-val* = 2.22; time: 1.21-1.28s) and left primary auditory cortices (Fig. 2E, parcel 2, main cluster I: *p* < .001, k = 12, *t-val* = - 3.70; time: 0.74-0.81s; II: *p* < .001, k = 12; *t-val* = 3.09; time: 0.87-0.95s; III: *p* < .001, k = 9; *t-val* = 2.90; time: 0.64-0.69s). With a reduced strength, similar clusters of activity have been observed for right (Fig. 2E, parcel 3, main cluster I: *p* < .001, k = 13, *t-val* = 3.08; time: 1.19-1.27s; II: *p* < .001, k = 12; *t-val* = 3.61; time: 0.89-0.96s) and left secondary auditory cortex and hippocampal areas (Fig. 2E, parcel 4, main cluster I: *p* < .001, k = 12, *t-val* = -2.97; time: 0.74-0.81s; II: *p* < .001, k = 10; *t-val* = 2.86; time: 0.87-0.93s) and cingulate (Fig. 2E, parcel 5, main cluster I: *p* < .001, k = 10, *t-val* = 3.03; time: 0.90-0.96s; II: *p* < .001, k = 9; *t-val* = - 2.29; time: 0.79-0.84s). Additional details on these contrasts are reported in **Tables S7**, and extensively depicted in Fig. S6 and S7.

### Modelling the brain activity underlying temporal pattern recognition

Once the difference between M and N was largely proved and detailly described, we focused on a further aim of the study, which was to mathematically characterize the dual simultaneous brain processing happening during recognition of the previously memorized temporal patterns. Thus, we computed another round of k-means functional clustering. This time, such analysis was performed only on the group-level main effect of M, to outline a functional parcellation based on the sole recognition of previously memorized patterns. As shown in Fig. 2F, the algorithm returned similar results compared to the previous round of k-means functional clustering, but better highlighted the parcels comprising brain areas implicated in memory and evaluative processes.

With regards to modelling, we hypothesized two different mathematical equations (one for each frequency band) that could describe the brain activity over our functionally defined parcels for global and local brain processes (Fig. 2D2, bottom).

Regarding delta (global processing of the pattern), we used a simple Gaussian function, described as follows:

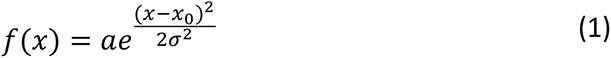

where *a* modulates the amplitude of the curve, *x_0_* shifts it over time and *σ* determines its width. This equation was fitted using a widely adopted non-linear least square approach, whose results are reported in Table 2 and **S8** and depicted in Fig. 2F (left column), S8 and S9. This procedure returned rather good results, highlighting key similarities and differences between the parcels timeseries. As reported in Table 2, the main functional parcels returned a similar peak amplitude (*a*). Conversely, the latencies of such peaks were highly different and shifted over time, as illustrated by parameter *x_0_*. Further, the width of the Gaussian function (indexed by *σ*) varied over the parcels. Indeed, lower-level brain areas such as right and left auditory cortices presented a reduced width compared to higher-level brain areas such as cingulate, insula, hippocampus, inferior temporal cortex and frontal operculum. This result may suggest that the transition from low- to high-order brain areas at the basis of the global processing of the pattern, is also reflected in a longer computation of the information operated by the brain.

**Table 2.** R^2^ and coefficients derived from non-linear least square fitting of the equations (1) and (3) on the brain activity underlying temporal pattern recognition. i refers to the five objects (musical tones) forming the temporal patterns. Here, the parcel IDs correspond to the ones reported in Fig. 2A1 and 2A2.

| 0.1 – 1 Hz |  |  |  |  | 2 - 8Hz |  |  |  |  |  |  |  |
| --- | --- | --- | --- | --- | --- | --- | --- | --- | --- | --- | --- | --- |
| Parcel # | R<sup>2</sup> | a | x<sub>0</sub> | σ | Parcel # | R<sup>2</sup><sub>i</sub> | i | a<sub>i</sub> | x<sub>0i</sub> | σ<sub>i</sub> | c<sub>i</sub> | φ<sub>i</sub> |
| 1 | 0,10 | 3,68 | 69,79 | 30,11 | 1 | 0,99 | 1 | -5,52 | -1,18 | 0,05 | -0,75 | 1,72 |
|  |  |  |  |  |  |  | 2 | 14,82 | 3,72 | 0,75 | 0,16 | 3,40 |
|  |  |  |  |  |  |  | 3 | -18,24 | 3,75 | 0,27 | 0,21 | -0,45 |
|  |  |  |  |  |  |  | 4 | -2,64 | 4,48 | 0,52 | 0,33 | -7,42 |
|  |  |  |  |  |  |  | 5 | 6,61 | 5,34 | 0,08 | 0,16 | -7,87 |
| 2 | 0,78 | 2,40 | 122,91 | 34,99 | 2 | 0,97 | 1 | -5,52 | -1,18 | 0,05 | -0,75 | 1,72 |
|  |  |  |  |  |  |  | 2 | 11,84 | 3,78 | 0,79 | 0,16 | 2,55 |
|  |  |  |  |  |  |  | 3 | -27,29 | 3,73 | 0,29 | 0,19 | 0,09 |
|  |  |  |  |  |  |  | 4 | -1,65 | 4,52 | 0,66 | 0,33 | -7,35 |
|  |  |  |  |  |  |  | 5 | 7,06 | 5,39 | 0,07 | 0,16 | -9,00 |
| 3 | 0,92 | 2,12 | 186,79 | 63,61 | 3 | 0,97 | 1 | -5,52 | -1,18 | 0,05 | -0,75 | 1,72 |
|  |  |  |  |  |  |  | 2 | 13,73 | 3,81 | 0,60 | 0,16 | 2,96 |
|  |  |  |  |  |  |  | 3 | -15,78 | 3,81 | 0,27 | 0,20 | 0,61 |
|  |  |  |  |  |  |  | 4 | -1,84 | 4,23 | 0,69 | 0,33 | -7,90 |
|  |  |  |  |  |  |  | 5 | 5,85 | 5,32 | 0,07 | 0,14 | -4,29 |
| 4 | 0,92 | 3,60 | 178,79 | 44,04 | 4 | 0,97 | 1 | -5,52 | -1,18 | 0,05 | -0,75 | 1,72 |
|  |  |  |  |  |  |  | 2 | 8,76 | 3,75 | 0,73 | 0,16 | 2,42 |
|  |  |  |  |  |  |  | 3 | -19,57 | 3,73 | 0,29 | 0,19 | 0,50 |
|  |  |  |  |  |  |  | 4 | -1,29 | 4,38 | 0,70 | 0,33 | -7,41 |
|  |  |  |  |  |  |  | 5 | 4,70 | 5,38 | 0,07 | 0,17 | -10,13 |
| 5 | 0,78 | 2,25 | 232,61 | 52,83 | 5 | 0,96 | 1 | -5,52 | -1,18 | 0,05 | -0,75 | 1,72 |
|  |  |  |  |  |  |  | 2 | 4,52 | 3,82 | 0,25 | 0,19 | 2,39 |
|  |  |  |  |  |  |  | 3 | -3,44 | 4,10 | 0,72 | 0,23 | -1,75 |
|  |  |  |  |  |  |  | 4 | -5,90 | 4,43 | 0,22 | 0,21 | 2,84 |
|  |  |  |  |  |  |  | 5 | 4,40 | 5,28 | 0,06 | 0,17 | -10,22 |

Conversely, with regards to theta (local processing of the objects forming the pattern), we hypothesized the following equation:

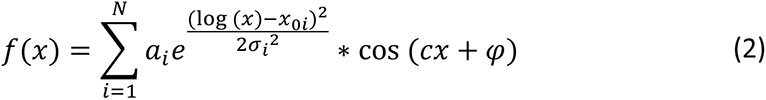

where *a*, *x_0_*, *σ* describes a Gaussian function, exactly as reported for equation (1). This new equation gives rise to a sinusoidal curve that modulates its amplitude on the basis of the associated Gaussian function. As usual, *c* refers to the angular frequency, while *φ* indicates the phase. Finally, *N* refers to the total number of objects forming the temporal pattern. This equation was hypothesized since it produces ‘wavelet-like’ timeseries, arguably describing the well-established series of components (peaks of activity in the timeseries, e.g. P50, N100, P300, N300 ^33^) generated by the brain in response to a sound. Indeed, such components have different latencies with respect to the sound onset and present opposite polarities (i.e. P50 and P300 are positive, while N100 and N300 are negative), giving rise to a wavelet-looking timeseries. Although well-established, it is not clear how these components relate to each other, especially when there are multiple brain sources involved and during complex cognitive processes such as recognition of temporal patterns. As done for equation (1), equation (2) was also fitted using the non-linear least square algorithm, returning good results (reported in Table 2 and **S8** and depicted in Fig. 2F, right column). However, in this case, the interpretation of the fitted parameters was more complicated since the brain responses to any two subsequent sounds was partially overlapping (i.e. the N300 component enhanced by the first sound occurred with a latency of approximately 320ms and overlapped with the P50 component arising after 50ms from the onset of the second sound). This fact partially altered the contour of the ‘wavelets’ and made the interpretation of the parameters less straightforward. Nevertheless, *x_0_* showed that the center of the ‘wavelets’ was progressively shifted over time following the onset of the sounds. Moreover, *a* indicated a trend of decreased absolute value over time, coherently with the reduced amplitude of the ‘wavelets’ occurring for the last sounds of the pattern.

Finally, Fig. 2G illustrates that while the ‘wavelet’ response to the first sound showed very similar activity over primary (*parcel i*) and secondary auditory cortices, insula, hippocampal areas (*parcel ii*) and cingulate (*parcel iii*), the peaks for the following sounds showed a different trend, especially in response to the third and fourth object of the pattern. In this case, secondary auditory cortices, insula, hippocampal areas and cingulate seemed to peak before the primary auditory cortex. However, this does not indicate a faster response of those areas as it could be thought at a first sight. Indeed, looking, for example, at the peaks around 0.5 seconds (first red square in Fig. 2G), the first peak (mainly occurring for secondary auditory cortices, insula, hippocampal areas and cingulate) corresponded to the P300 component to the second sound of the pattern, while the second peak (mainly occurring for primary auditory cortex) was the P50 to the third sound. An analogous phenomenon happened for the following objects of the pattern (as outlined by the other red squares). This shows that while the contribution of the primary auditory cortex was stronger for the first components (i.e. P50 and N100), which indexed lower-level processes, later components such as P300 were mainly generated by higher order areas such as secondary auditory cortices, insula, hippocampal areas and cingulate.

## Discussion

In this study, combining MEG and MRI, we discovered and mathematically modelled an inedited dual simultaneous processing responsible for brain recognition of temporal patterns. Indeed, on the one hand the single objects forming the pattern were independently elaborated by a rapid (theta band), oscillatory, local processing driven by sensorial cortices.

On the other hand, the combination of the single objects into a meaningful superordinate pattern depended on a simultaneous global, slow (delta band) processing involving a widespread network of sequentially active high-order brain areas.

Our findings revealed that the dual simultaneous processing required by the brain to recognize temporal patterns involved a widespread network of brain areas largely related to memory, attention, audition, and decision-making. Such brain areas were hippocampus ^34^, cingulate gyrus ^35^, inferior temporal cortex ^36^, frontal operculum ^37,38^, insula ^39^, and primary and secondary auditory cortex ^40^. Notably, both processes (global and local) involved approximately the same brain regions but depended on different frequencies of the neural signal. Furthermore, as conceivable, the local processing relied mainly on sensorial cortices (e.g. auditory cortex), while the global processing presented a wider recruitment of higher-order brain areas such as cingulate, inferior temporal cortex and hippocampus.

Strikingly, temporal pattern recognition occurred through a cascade of progressively slower events rewiring a chain of low- to high-order brain regions, as formally described by our modelling. This evidence, observed for delta band, may indicate that the brain progressively constructs a meaningful understanding of the unfolding temporal pattern by recruiting a hierarchical pathway of subsequently active regions. Conversely, theta band activity showed a complementary profile. Indeed, its activity peaked slightly after each object of the temporal sequence. Such evidence suggests that, while delta band may be implicated in achieving a comprehensive understanding of the whole pattern (global processing), theta may elaborate independently its objects (local processing). Notably, previous research described global and local processing mainly in terms of different locations of the neural signal (i.e. primary sensorial cortices preceded higher-order brain areas in the elaboration of incoming stimuli ^16,17^). Conversely, in our study we showed that the same brain regions operated these two processes (global and local) at the same time, using two concurrent frequency bands.

Further, previous research on memory for music and auditory sequences showed the involvement of large brain networks ^23–26^, but did not reveal any dual simultaneous processing nor detailly described the dynamical, rapid change of the brain areas’ activity in relation to the development of the auditory stimuli. Moreover, the majority of such studies employed fMRI, a powerful tool which returns excellent spatial resolution but lacks temporal accuracy ^41^. On another note, recent studies on musical memory benefitting from the excellent temporal resolution of MEG focused on different features of memory, mainly investigating working memory paradigms and retention of musical information ^42,43^. In conclusion, on the one hand our study highly confirmed and refined classic results on auditory and musical memory. On the other hand, it proposed a novel mechanism used by the brain to extract meaning from temporal sequences, shedding new light on the brain strategies to process, and become aware of the complexity of the external environment.

Another crucial evidence emerged from our study relates to the differential strength of the brain signal observed for the two frequency bands in relation to our experimental conditions (M and N). Indeed, while delta band presented a stronger power for the memorized patterns, theta showed greater responses for the novel ones. This finding may be seen in light of the predictive coding theory ^44,45^, which posits that the brain is constantly updating internal models to predict environmental information. Here, when the brain is recognizing the temporal patterns (e.g. around tones number two and three of our sequences), it might formulate better predictions of the upcoming, previously memorized, objects completing the patterns. Thus, such objects would require a lower local processing, as we observed experimentally. Interestingly, although mainly localized in primary auditory cortex, the neural sources of theta band activity were also placed in hippocampal areas, secondary auditory cortex, insula, and cingulate. As previously mentioned, this evidence suggests that roughly the same brain regions generated two simultaneous frequency bands characterized by a very different functional profile, indexing the local and global processing of the temporal pattern. On top of this, with regards to local processing, our results show that the elaboration of each sound gave rise to a wavelet-like timeseries with three main peaks (components). Here, the lower-level elaboration of the sounds indexed by the first components (i.e. P50 and N100 ^46^) originated mainly in the primary auditory cortex. Conversely, later components such as P300 ^46^ were generated especially by higher order areas such as secondary auditory cortices, insula, hippocampal regions and cingulate. Remarkably, such phenomenon became more evident following the unfolding of the temporal pattern, suggesting that a progressively more refined elaboration of the single objects is essential for the brain to comprehend the meaning of the whole temporal pattern.

Finally, our findings related and expanded concepts of the notorious two-stream hypothesis of the brain ^47,48^. Such conceptualization proposed two main pathways for high-order elaboration of visual and auditory information. On the one hand, the ventral stream leads from sensorial areas (e.g. visual and auditory cortices) to the medial temporal lobe, processing features mainly associated to object recognition ^48,49^. On the other hand, the dorsal stream brings information from sensory cortices to the parietal lobe, elaborating spatial features of the stimuli ^50^. Coherently with such hypothesis, our results highlighted several brain regions of the ventral stream that are implicated in recognition processes, such as hippocampal areas, frontal operculum, and inferior temporal cortex. Remarkably, our results further expanded previous knowledge on the two-stream hypothesis by providing at least three crucial remarks. The brain recognition of temporal patterns presented unique spatial-temporal features which were not shared with the identification of single elements or synchronous patterns (i). In addition to the brain regions involved in the two-stream hypothesis, our findings showed the privileged role of cingulate gyrus to achieve temporal pattern recognition (ii). Finally, the recognition of sequential patterns unfolding over time involved a dual simultaneous processing of the same objects, which the brain interpreted concurrently as individual pieces of information (local processing) and elementary parts of a larger reality (global processing) (iii).

In conclusion, in our study we achieved a rather profound understanding of the brain mechanisms underlying conscious recognition of temporal patterns. Indeed, we discovered and mathematically modelled a rapid (theta band), oscillatory, local processing driven by sensorial cortices responsible for the elaboration of the single objects (sounds) forming the pattern. Additionally, our findings suggested that the combination of such single objects into a meaningful superordinate pattern depended on a simultaneous global, slow (delta band) processing involving a widespread network of sequentially active high-order brain areas. By showing that nearly the same brain regions operated two processes at the same time using two concurrent frequency bands, our results unravelled the brain mechanisms underlying temporal pattern recognition and proposed a novel understanding of the strategies adopted by the brain to elaborate the complexity of the external environment.

## Methods

### Participants

The study comprised 70 volunteers: 36 males and 34 females (age range: 18 – 42 years old, mean age: 25.06 ± 4.11 years). All participants were healthy and reported no previous or current alcohol and drug abuse. Moreover, they were not under any kind of medication, declared that they did not have any previous neurological or psychiatric disorder, and reported to have normal hearing. Furthermore, their economic, educational and social status was homogeneous.

All the experimental procedures were carried out complying with the Declaration of Helsinki – Ethical Principles for Medical Research and were approved by the Ethics Committee of the Central Denmark Region (De Videnskabsetiske Komitéer for Region Midtjylland) (Ref 1-10-72-411-17).

### Experimental design and stimuli

To detect the brain signature of temporal pattern recognition, we used an old/new paradigm ^51^ auditory recognition task during magnetoencephalography (MEG) recording. First, participants listened to four repetitions of a MIDI homo-rhythmic version of the right-hand part of the whole prelude BWV 847 in C minor composed by J.S. Bach (total duration of about 10 minutes). Second, they were presented with 80 brief musical excerpts lasting 1250 ms each and were asked to state whether each excerpt belonged to the prelude by Bach (‘memorized’ pattern (M), old) or it was a variation of the original patterns (‘novel’ pattern (N), new) (Fig. 1A). Forty excerpts were taken from the Bach’s piece and 40 were novel. In the following analysis we used only the correctly recognized trials (mean correct M: 78.15 ± 13.56 %, mean reaction times (RT): 1871 ± 209 ms; mean correct N: 81.43 ± 14.12 %, mean RT: 1915 ± 135 ms). Both prelude and excerpts were created by using Finale (MakeMusic, Boulder, CO) and presented with Presentation software (Neurobehavioural Systems, Berkeley, CA). After the acquisition of the MEG data, in the same or another day, participants’ brain structural images were acquired by using magnetic resonance imaging (MRI).

### Data acquisition

We acquired both MRI and MEG data in two independent sessions. The MEG data was acquired by employing an Elekta Neuromag TRIUX system (Elekta Neuromag, Helsinki, Finland) equipped with 306 channels. The machine was positioned in a magnetically shielded room at Aarhus University Hospital, Denmark. Data was recorded at a sampling rate of 1000 Hz with an analogue filtering of 0.1–330 Hz. Prior to the measurements, we accommodated the sound volume at 50 dB above the minimum hearing threshold of each participant. Moreover, by utilizing a three-dimensional digitizer (Polhemus Fastrak, Colchester, VT, USA), we registered the participant’s head shape and the position of four headcoils, with respect to three anatomical landmarks (nasion, and left and right preauricular locations).

The location of the headcoils was registered during the entire recording by using a continuous head position identification (cHPI), allowing us to track the exact head location within the MEG scanner at each time-point. We utilized this data to perform an accurate movement correction at a later stage of the data analysis.

The recorded MRI data corresponded to the structural T1. The acquisition parameters for the scan are reported as follows: voxel size = 1.0 x 1.0 x 1.0 mm (or 1.0 mm^3^); reconstructed matrix size 256×256; echo time (TE) of 2.96 ms and repetition time (TR) of 5000 ms and a bandwidth of 240 Hz/Px. At a later stage of the analysis, each individual T1-weighted MR scan was co-registered to the standard MNI brain template through an affine transformation and then referenced to the MEG sensors space by using the Polhemus head shape data and the three fiducial points measured during the MEG session.

### Data pre-processing

The raw MEG sensor data (204 planar gradiometers and 102 magnetometers) was pre-processed by MaxFilter ^52^ for attenuating the interference originated outside the scalp by applying signal space separation. Within the same session, Maxfilter also adjusted the signal for head movement and down-sampled it from 1000 Hz to 250 Hz.

The data was converted into the SPM format and further analyzed in Matlab (MathWorks, Natick, Massachusetts, United States of America) by using OSL, a freely available toolbox that relies on a combination of FSL ^53^, SPM ^54^ and Fieldtrip ^55^, as well as in-house-built functions. The data was then high-pass filtered (0.1 Hz threshold) to remove frequencies that were too low for being originated by the brain. A notch filter (48-52 Hz) was applied to correct for possible interference of the electric current. The data was further down-sampled to 150 Hz and few parts of the data, altered by large artifacts, were removed after visual inspection. Then, to discard the interference of eyeblinks and heart-beat artefacts from the brain data, independent component analysis (ICA) was used to decompose the original signal in independent components. Then, the components that picked up eyeblink and heart-beat activities were first isolated and then discarded. The signal was rebuilt by using the remaining components ^56^ and then epoched in 80 trials (one for each musical excerpt) lasting 3500 ms each (with 100ms of pre-stimulus time that was used for baseline correction) (Fig. 1B).

### Univariate tests and Monte-Carlo simulations over MEG sensors

Although our primary focus was on the MEG source reconstructed brain data, a first analysis on MEG sensors data was computed, coherently with state-of-the-art recommendation about best practice in MEG analysis ^57^.

Thus, according to a large number of MEG and electroencephalography (EEG) task-related studies ^57^, we averaged the trials over conditions, obtaining two final mean trials, for M and N, respectively. Then, we combined each pair of planar gradiometers by root sum square. Afterwards, we performed a t-test for each time-point in the time-range 0 – 2.500 seconds and each combined planar gradiometer, contrasting M vs N. To correct for multiple comparisons, we computed Monte-Carlo simulations (MCS) ^58^ with 1000 permutations on the clusters of significant results emerged from the t-tests. We considered significant the original clusters that had a size bigger than the 99.9% maximum cluster sizes of the permuted data. Additional details on this widely used procedure can be found in Bonetti and colleagues ^29,30^. This analysis returned a large and robust difference between experimental conditions. Moreover, the brain activity recorded over the MEG channels forming the significant cluster outputted by the MCS analysis outlined a timeseries which presented two main frequency components. As shown in Fig. S1A, the faster frequency component peaked after the presentation of each of the objects forming the pattern, while the slower frequency component accompanied the whole pattern, peaking at its end. This evidence was further supported by the computation of complex Morlet wavelet transform ^59^ on all MEG sensor data, which highlighted the main contribution of 1Hz and 4Hz to the MEG signal recorded during the task (Fig. S1B). Thus, our following analyses in source reconstructed space focused on two frequency bands defined around those main frequencies: 0.1-1Hz and 2-8Hz. These bands roughly corresponded to the well-known brain waves called delta and theta, respectively ^60^, and were also coherent with results reported by Bonetti et al. ^29^. Importantly, we hypothesized that they indexed the two main processes involved in our experimental task: processing of single objects forming the temporal pattern (*i - local processing*) and recognizing the temporal pattern as a comprehensive superordinate object (*ii - global processing*).

### Source reconstruction

MEG is a powerful tool to record whole-brain activity with excellent temporal resolution. However, the investigation of neural activity also requires spatial parameters. To achieve a reasonably accurate information about the brain sources that generated the MEG signal, an inverse problem must be solved. Indeed, from MEG recording we know the power of the neural signal outside the head, but we do not know which brain sources generated it. Moreover, we possess only 102 triplets of MEG sensors, while the active brain sources that could be distinctly recorded by the MEG are much more numerous. To solve this problem, state-of-the-art source reconstruction methods have been used (Fig. 1C and Fig. 2A) ^61,62^. Importantly, the source reconstruction algorithm has been computed independently for the two frequency bands involved in the study (0.1 – 1 Hz and 2 – 8 Hz (Fig. 1C). Specifically, the following steps were implemented. First, the continuous data (before the epoching) was band-pass filtered into the two frequency bands. Second, the filtered data (independently for the two bands) was epoched. Third, the epoched data was submitted to the source reconstruction algorithm described below. Such algorithm involves two subsequent steps: (i) designing a forward model; (ii) computing the inverse solution. The forward model is a theoretical model which considers each brain source as an active dipole and describes how the unitary strength of such dipole would be reflected over all MEG sensors (in our case we utilized both magnetometers and planar gradiometers) ^62^. Here, we employed an 8-mm grid which returned 3559 dipole locations (voxels) within the whole brain. After co-registering individual structural T1 data with fiducials (information about head landmarks), the forward model was computed by adopting a widely used method called “Single Shell”, presented in details by ^63^. The output of such computation, also referred to as leadfield model, was stored in matrix *L* (sources x MEG channels). In the few cases where structural T1 was not available, we performed the leadfield computation using a template (MNI152-T1 with 8-mm spatial resolution).

The second step of the source reconstruction is to compute the inverse solution (i.e. to estimate the generators of the neural signal on the basis of the brain activity recorded with MEG). In our study, we chose the beamforming, which is one of the most popular and effective algorithms available in the field ^61,62^. This procedure uses a different set of weights sequentially applied to the source locations for isolating the contribution of each source to the activity recorded by the MEG channels for each time-point ^61,64^. On a more technical level, the inverse solution based on beamforming can be described by the following main steps.

First, the data recorded by MEG sensors (*B*) at time *t*, can be described by the following equation (1):

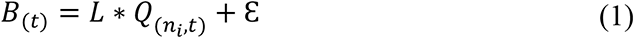

Where *L* is the above-described leadfield model, *Q* is the dipole matrix carrying the activity of each active dipole (*q*) over time and *Ɛ* is noise (see Huang and colleagues ^65^ for more details). Thus, to solve the inverse problem, we have to compute *Q*. Using the beamforming, such procedure revolves around the computation of weights that are applied to the MEG sensors at each time-point, as shown for the single dipole *q* in equation (2):

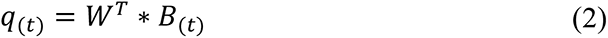

Indeed, to gain *q*, the weights *W* should be computed (the subscript *T* refers to transpose matrix). To do so, the beamforming relies on the matrix multiplication between *L* and the covariance matrix between MEG sensors (*C*), computed on the concatenated experimental trials. Specifically, for each brain source *n*, the weights *W_n_* are computed as follows:

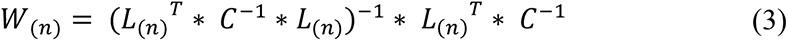

To be noted, the computation of the leadfield model was done for the three main orientations of each brain source (dipole), according to Nolte ^63^. However, before computing the weights, the orientations have been reduced to one by using the singular value decomposition algorithm on the matrix multiplication reported in equation (4). This procedure is widely adopted to simplify the beamforming output ^65,66^.

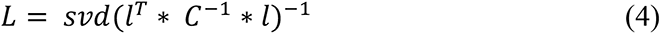

Here, *l* represents the leadfield model with the three orientations, while L the resolved one-orientation model that was used in (3).

Finally, with regards to the coding implementation of such algorithms, we have used Matlab toolboxes such as OSL, FieldTrip, SPM (functions for MEEG preprocessing and SPM beamforming toolbox) and FSL. Moreover, those codes were complemented by in-house-built scripts and functions.

Importantly, as already highlighted above, the analyses reported in the following paragraphs have been computed independently for the two frequency bands considered in the study.

### Brain activity for each element of the temporal sequence

First, we wanted to detect the brain activity underlying each object of our temporal patterns (Fig. 1D, Fig. S2, Table 1 and **Table S1**). Here, we computed the absolute value of the reconstructed time-series since we were interested in the absolute strength of the signal, and we wanted to avoid the sign ambiguity introduced by source reconstruction procedures.

To perform first-level analysis for each participant, we employed general linear models (GLMs). Such models were computed on the source reconstructed data for each time-point and brain source ^31^. The GLMs returned the main effect (contrasts of parameters estimate (COPEs)) of M and N as well as their contrast. These results were submitted to a second-level analysis, employing one-sample t-tests with spatially smoothed variance obtained with a Gaussian kernel (full-width at half-maximum: 50 mm) ^65^.

Here, we were interested in observing the different brain activity underlying recognition of M vs N temporal sequence, independently for each frequency band and object forming the sequence (musical tone). Thus, we computed 10 (five tones x two frequency bands) cluster-based Monte-Carlo simulations (MCS) on the second level (group-level) analysis results averaged over the five time-windows corresponding to the duration of the musical tones. The MCS analysis comprises 1000 permutations and a cluster forming threshold of *p* < .05 (from the second-level t-tests). Specifically, the MCS test consisted of detecting the spatial clusters of significant values in the original data. Then, such data was permuted and the spatial clusters of the permuted significant values were detected. This procedure was computed several times (e.g. 1000) and gave rise to a reference distribution of cluster sizes detected for each permutation. Finally, the original cluster sizes were compared to the reference distribution. The original clusters were considered significant if the cluster sizes of the permuted data were bigger than the original cluster sizes less times than the MCS *α* level. In this case, since we computed the analysis 10 times, we corrected for multiple comparisons by dividing the standard MCS *α* level (= .05) by 10, resulting in an updated MCS *α* = .005 (i.e. original clusters were significant if their sizes were larger than the 99.5% of the permuted cluster sizes).

### K-means functional clustering

Contrasting the brain activity in response to each element (musical tone) forming the temporal sequence is an effective procedure to obtain a general understanding of the brain functioning underlying the discrete processing of the sequence.

However, this strategy does not fully benefit of the excellent temporal resolution of the MEG data and underestimate brain processes that may happen in between two or more objects of the sequence. Furthermore, computing contrasts for each time-point and brain source returns a large amount of data which is partially redundant and sometimes not straightforward to understand and ideal to mathematically model. Indeed, several brain sources are highly correlated because of both biological reasons involving large populations of neurons generating the signal and artificial source leakage introduced during the source reconstruction ^32^. Thus, defining a functionally based parcellation of the brain may be of great importance when aiming to synthesize and mathematically describe the spatial extent of the active brain sources as well as their activity over time.

To overcome these issues, we adopted a so-called k-means functional clustering, consisting of a series of k-means clustering algorithms ^67^ performed on functional and spatial information of each of the reconstructed brain sources (voxels) timeseries.

Specifically, as a first step this algorithm computed a clustering on basic functional parameters such as peak values and the corresponding indices of the voxels timeseries. We refer to this step as *functional clustering*. This procedure returned a set of independent parcels grouped according to the functional profiles of the brain voxels. Indeed, such parcels could either contain voxels that peaked approximately at the same time (Fig. 2B, left) or with similar absolute strength (Fig. 2B, right). As conceivable, clustering on the maximum timeseries indices is suggested when the brain activity is localized in different regions at different times. Conversely, when the activity is highly correlated over most of the brain voxels, clustering should be done on maximum timeseries values and would help to identify the core generators of the neural signal. In this study, delta (global processing of the pattern) presented different peaks of activity shifted over time and thus was clustered considering the time-indices of such peaks. Differently, theta (local processing of the pattern) showed very correlated activity and was therefore clustered using the absolute values of such peak activity. As widely done in clustering analysis ^68^, also in our case it was beneficial to compute the clustering algorithm on a sequential set of *k* clusters (from *k* = 2 to *k* = 20). Then, the best clustering solution was decided on the basis of well-known evaluation strategies (heuristics) such as the elbow method/rule ^69^ and the silhouette coefficient ^70^. The elbow method consists in plotting the sum of squared errors (SSE) of the elements belonging to the clusters with respect to the cluster’s centroids, as a function of the progressively more numerous cluster solutions. Then, the method suggests to visually identify the “elbow” of the curve as the number of clusters to use. The silhouette coefficient is a value (ranging from -1 to +1) showing the similarity of an element with its own cluster (cohesion) when compared to other clusters (separation). A high silhouette coefficient value indicates that the element is well matched to its own cluster and poorly to the neighboring clusters. As conceivable, if most of the elements present a high value, the clustering configuration is appropriate. We reported a further graphical example of this method in Fig. S3.

Once the best functional clustering solution is decided, a second clustering with regards to spatial information should be computed (*spatial clustering*, Fig. 2C). Indeed, brain activity is mainly described by two parameters, spatial locations, and variation over time. Clustering the original brain voxels into distinct functional parcels may return large parcels involving a network of spatially separated brain areas that are e.g. active at the same time. Thus, to define a better parcellation, it is beneficial to conduct clustering analysis also on the spatial coordinates of each of the functional parcels. In our study, we considered the three-dimensional spatial coordinates (in MNI space) of the voxels forming each of the functional parcels. This clustering computation was performed for a sequential set of *k* clusters solutions (from *k* = 2 to *k* = 10), for one parcel at a time. As for the functional clustering, we evaluated the best solution for the spatial clustering by using the elbow rule and the silhouette coefficient. The k-means functional clustering was complete once this procedure was performed on all functional parcels, suggesting an effective parcellation for the experimental task based on both functional and spatial information (examples are reported in Fig. S4**, Table S2** and **S3** for 0.1-1 Hz and Fig. S5, **Table S4**, **S5** and **S6** for 2-8 Hz). As a last step, the timeseries of the brain voxels belonging to each parcel were averaged together to provide a final timeseries for the parcel. As follows, we provide a few conceptual remarks related to this algorithm and to the current study that should be highlighted.

First, the k-means functional clustering has to be computed on source reconstructed brain data. However, such data can be either the timeseries outputted by the source reconstruction or the timeseries of the statistics computed on the source reconstruction. Moreover, the algorithm can be computed independently for each participant or on the group level statistics. Further, the brain data in input can either be the broadband data or the data reconstructed in selected frequency bands. Moreover, in the likely case of having more than one experimental condition, as conceivable, the algorithm can be computed on each condition independently or on the aggregated (e.g. averaged) conditions. The best procedure cannot be defined a priori for every study and highly depends on the specific aims of the project. In our case, since the main aim of the algorithm was to define functional parcellations with timeseries that best represented the brain functioning among the whole population, we decided to work on the group level statistics. With regards to experimental conditions, we have computed different runs of the k-means functional clustering. Indeed, to statistically compare the timeseries of each parcel for the two experimental conditions of our task, we performed the clustering algorithm on the main effects of the two conditions averaged together. Instead, when aiming to mathematically describe the timeseries of each parcel for a specific experimental condition (e.g. M), the clustering algorithm has been computed on such condition only. Further, in relation to the frequency bands, we performed one computation of the clustering algorithm for each of the two frequency bands involved in our study. In the case of 0.1-1 Hz, we worked with absolute values of the reconstructed brain sources timeseries since they did not present any complete cycle of the oscillation, considering their absolute strength as sufficient. Conversely, when dealing with 2-8 Hz, the timeseries presented several complete oscillations and thus computing their absolute values would lead to lose important information. We resolved the sign ambiguity introduced by the source reconstruction by referencing the sign of the timeseries to the well-known negative polarity of the N100 emerged in response to the first tone of the pattern. Then, we computed the statistics and the subsequent k-means functional clustering on the timeseries which maintained their original double polarity.

Second, when dealing with real data, an “ideal” clustering solution may often not be existent, and the elbow method and silhouette coefficients may return slightly contradictory results and controversial conclusions. This is a quite usual limitation of clustering algorithms that, however, should not be necessarily interpreted as a threatening issue. Indeed, for instance, if the elbow method and silhouette coefficients indicate as the best solutions a series e.g. three subsequent *ks* but without clearly stating one single *k*, it is reasonable to expect very similar results among the three different *k* solutions. Thus, although this would suggest that an ideal *k* is probably not existent, it should be noted that any different choice of the suggested *ks* should not lead to a huge affection or misinterpretation of the final results. On the contrary, stating that an “ideal” solution may often not exist does not mean that clustering algorithms will invariably return a valid output. Indeed, such techniques are designed to always provide results, even in the cases where there are no reasons for clustering the data. With regards to the brain, extremely poor clustering solutions would be achieved when brain sources are all very similar in terms of functional and spatial profile. This should not happen if the data is acquired with properly designed experimental tasks, but it should always be born in mind as a realistic possibility. Importantly, in such a case, the elbow method and silhouette coefficients would not return any clear indication regarding the ideal number *k* of clusters and the clustering algorithm would therefore be highly not recommended.

Third, it is important to state that the clustering procedure that we described here has been developed for task-related MEG data and would not properly work for resting state scenarios where other algorithms such as principal component analysis (PCA) ^71^ may be more appropriate.

Fourth, to increase the strength of the clustering algorithm, it may be beneficial to zero the activity of few poorly active brain sources timeseries before computing the functional k-means clustering. This action would help the clustering procedure and may provide some beneficial effects for the definition of the functional parcellation.

### Contrasts over time for each parcel

Here, the k-means functional clustering was performed on the group-level main effects of M and N averaged together. Then, to obtain the main effect of M and N for each parcel and participant, we averaged the first-level main effect of M and N (from the GLMs) over the brain voxels belonging to each of the functional parcels. This resulted in a new timeseries for each participant, functional parcel, and experimental condition (M and N). Such timeseries were submitted to univariate contrasts (M vs N; Fig. 2D1, methods, and Fig. 2E, S6 and S7, results). Specifically, for each parcel and time-point, we computed one two-sample t-test (threshold *p* < .05) contrasting the main effect of M vs N. Then, we corrected for multiple comparison by using a two-dimensional MCS approach with 1000 permutations. First, temporal clusters of significant results emerged from the t-tests were individuated. Second, significant results were permuted along the time dimension and clusters of such permuted results were identified. This procedure was computed 1000 times, giving rise to a reference distribution of cluster sizes of permuted results. Such distribution has been compared with the clusters size of the original data. Significant clusters in the original data were the ones whose size was bigger than the 99.9% of permuted cluster sizes (MCS *p* < .001). More details on this widely adopted statistical procedure can be found in Bonetti and colleagues ^29,72^. As done for the other analyses, such operation was observed for both frequency bands investigated in the study (Fig. 2 and **Table S7**).

### Curve fitting

Besides comparing our two experimental conditions, a main aim of our study was to mathematically describe the timeseries of the brain activity associated to recognition of temporal patterns (Fig. 2D2, methods, and Fig. 2F, S8 and S9, results). Indeed, we believe that to properly understand a scientific phenomenon, a mathematical description of such phenomenon should be provided, as commonly done in many branches of science. In addition, this procedure is a first essential step to develop future generative models that will not only describe the brain activity but simulate and perturbate its nature.

Thus, we computed another round of k-means functional clustering. This time, such analysis was performed only on the group-level main effect of M, to outline a functional parcellation based on the sole recognition of previously memorized patterns. Once again, this procedure was computed independently for the two frequency bands considered in the study.

Then, to describe the dual simultaneous brain processing happening during recognition of temporal patterns, we hypothesized two different mathematical equations (one for each frequency band).

Regarding the slower frequency band included in our study (0.1-1Hz), we used a simple Gaussian function, described as follows:

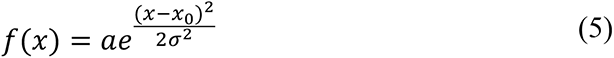

where *a* modulates the amplitude of the curve, *x_0_* shifts it over time and *σ* determines its width. In a few cases, we employed a modified version of the equation (5), which is basically a summation of three Gaussian functions shifted over time, as described as follows:

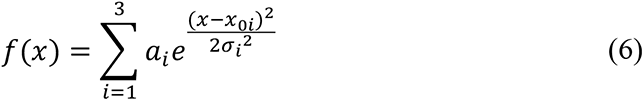

Arguably, this frequency indexed the recognition process of the full temporal pattern (global processing), as suggested by the brain areas involved and by the timing of their activations.

Conversely, with regards to the second frequency described in our study (2-8Hz) which supposedly reflected the brain processing of each object of the temporal sequence (local processing), we hypothesized the following equation:

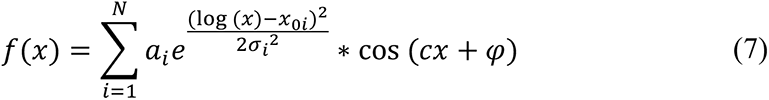

where *a*, *x_0_*, *σ* describes a Gaussian function, exactly as reported for equation (5) and equation (6). However, in this case, the Gaussian function can be ‘skewed’ by *log(x)* and is multiplied by a cosine function. This new equation gives rise to a sinusoidal curve that highly modulates its amplitude based on the associated Gaussian function. As usual, the parameter *c* refers to the angular frequency, while *φ* indicates the phase. Finally, *N* refers to the total number of objects (musical tone) forming the temporal pattern.

In all cases, the best parameters were fitted using the Python function curve_fit, which employs the widely adopted non-linear least squares method ^73^. Table 2 and **S8** reports the results of such analysis. In the few cases where no values are reported, the fitting of the equations were not possible since the timeseries referred to brain areas that were essentially not activate during our experimental tasks.

## Data availability

The codes are available at the following link: https://github.com/leonardob92/LBPD-1.0.git, while the multimodal neuroimaging data related to the experiment are available upon reasonable request.

## Acknowledgements

We thank Riccardo Proietti, Giulio Carraturo, Mick Holt, Holger Friis for their assistance in the neuroscientific experiment. We also thank the psychologist Tina Birgitte Wisbech Carstensen for her help with the administration of psychological tests and questionnaires and Francesco Carlomagno for his assistance with the display items of the paper. Furthermore, we thank the Linacre College in Oxford for its support to the author Leonardo Bonetti.

The Center for Music in the Brain (MIB) is funded by the Danish National Research Foundation (project number DNRF117).

LB is supported by the Carlsberg Foundation (CF20-0239) and by the Center for Music in the Brain.

MLK is supported by the ERC Consolidator Grant: CAREGIVING (n. 615539), Center for Music in the Brain, and Centre for Eudaimonia and Human Flourishing funded by the Pettit and Carlsberg Foundations.

GD is supported by the Spanish Research Project PSI2016-75688-P (AEI/FEDER, EU), by the European Union’s Horizon 2020 Research and Innovation Programme under grant agreements n. 720270 (HBP SGA1) and n. 785907 (HBP SGA2), and by the Catalan AGAUR Programme 2017 SGR 1545.

Additionally, we thank the University of Bologna for the economic support provided to the author Giulia Donati and the student assistants Riccardo Proietti and Giulio Carraturo and the Italian section *of Mensa: The* International High IQ Society for the economic support provided to Francesco Carlomagno.

## Author contributions

Conceptualization: LB, EB, MLK, PV; Methodology: LB, MLK, DP, GDE; Software: LB; Analysis: LB; Investigation: LB, GDO; Resources: MLK, PV, EB, LB; Data curation: LB; Writing - Original draft: LB; Writing – Review & editing: LB, SEPB, EB, DP, GDE, GDO, PV; Visualization: LB, SEPB; Supervision: MLK, PV, DP, EB; Project administration: LB, MLK, PV, EB; Funding acquisition: LB, PV, MLK.

## SUPPLEMENTARY MATERIALS

As follows, supplementary materials related to this study and organized as supplementary figures (i) and tables (ii). In the cases when the supplementary tables were too large to be conveniently reported in the current document, they have been reported in Excel files that can be found at the following link: https://www.dropbox.com/sh/sax1yzjqn897hxm/AAC8hWFE8IcyJgRCrJ2gu-bNa?dl=0)

## SUPPLEMENTARY FIGURES

**Fig. S1.**
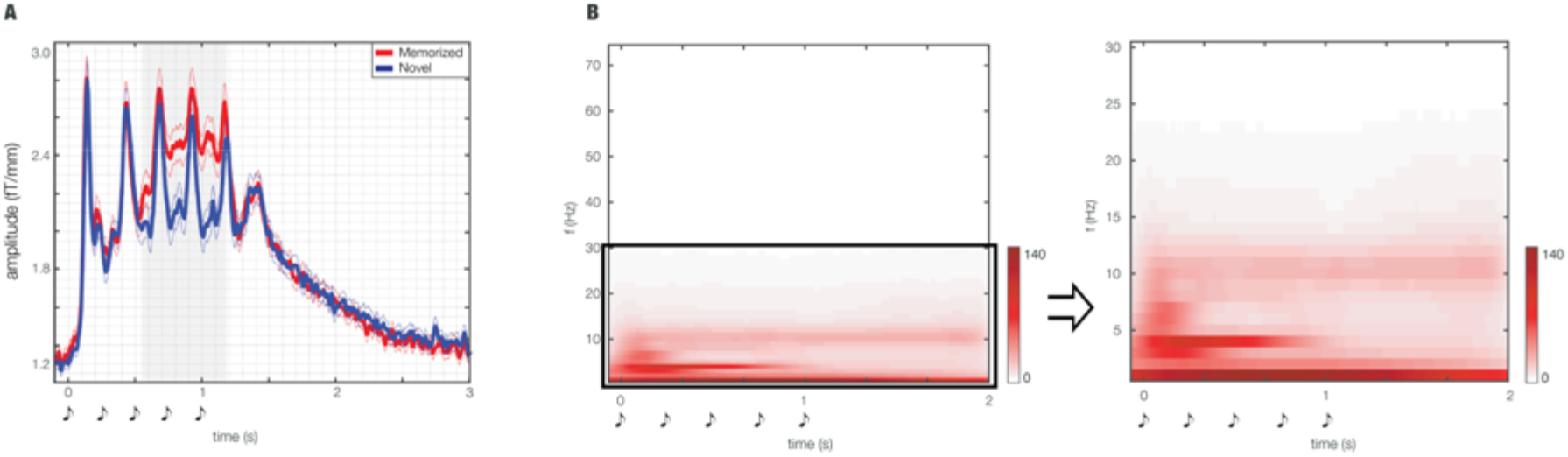
MEG sensors waveform and power spectra. The left plot shows the significantly different brain activity during recognition of ‘memorized’ vs ‘novel’ temporal sequences. The waveforms represent the average over the combined planar gradiometers forming the significant cluster emerged from the analysis, while the grey area illustrates the temporal extent of such significant difference. The complementary two plots show the power spectra computed by using complex Morlet wavelet transform on all MEG channels. The two plots illustrate the power spectra computed for progressively narrower bands. Together with the waveforms, these plots highlight the main contribution of the two frequency bands analyzed in the study: 0.1-1 Hz and 2-8 Hz (roughly corresponding to the well-known brain waves named delta and theta, respectively).

**Fig. S2.**
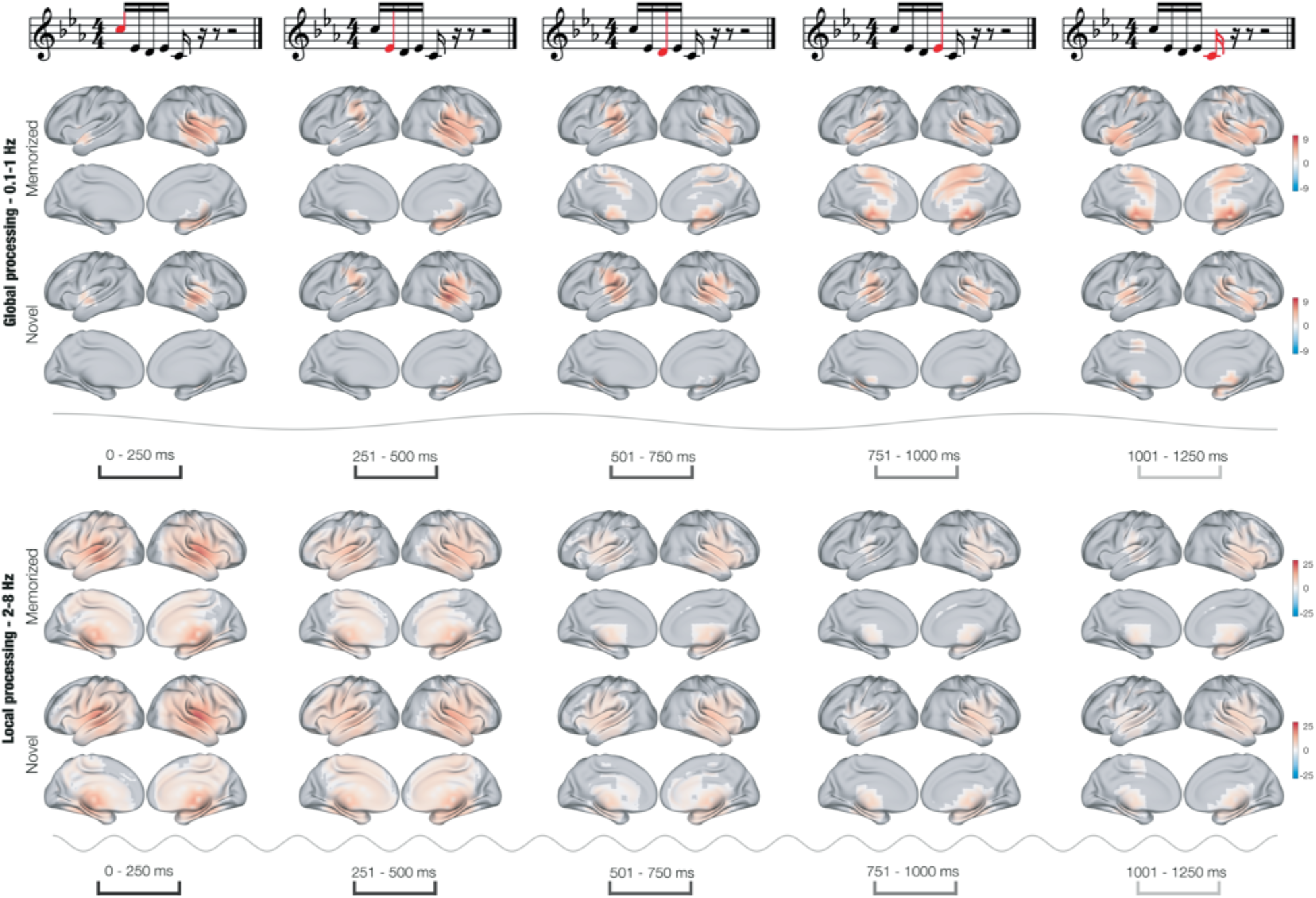
Brain activity underlying the single objects of the temporal patterns. Significant clusters of brain activity reconstructed in the time-windows corresponding to the five objects of the temporal patterns (as illustrated in the first row by the red tones). The brain activity shows the main effects for our experimental conditions (‘memorized’ and ‘novel’) and frequency bands (0.1-1Hz and 2-8Hz). The colorbars indicate one-sample t-values computed for each spatial location and time-point and then corrected with cluster-based permutation tests.

**Fig. S3.**
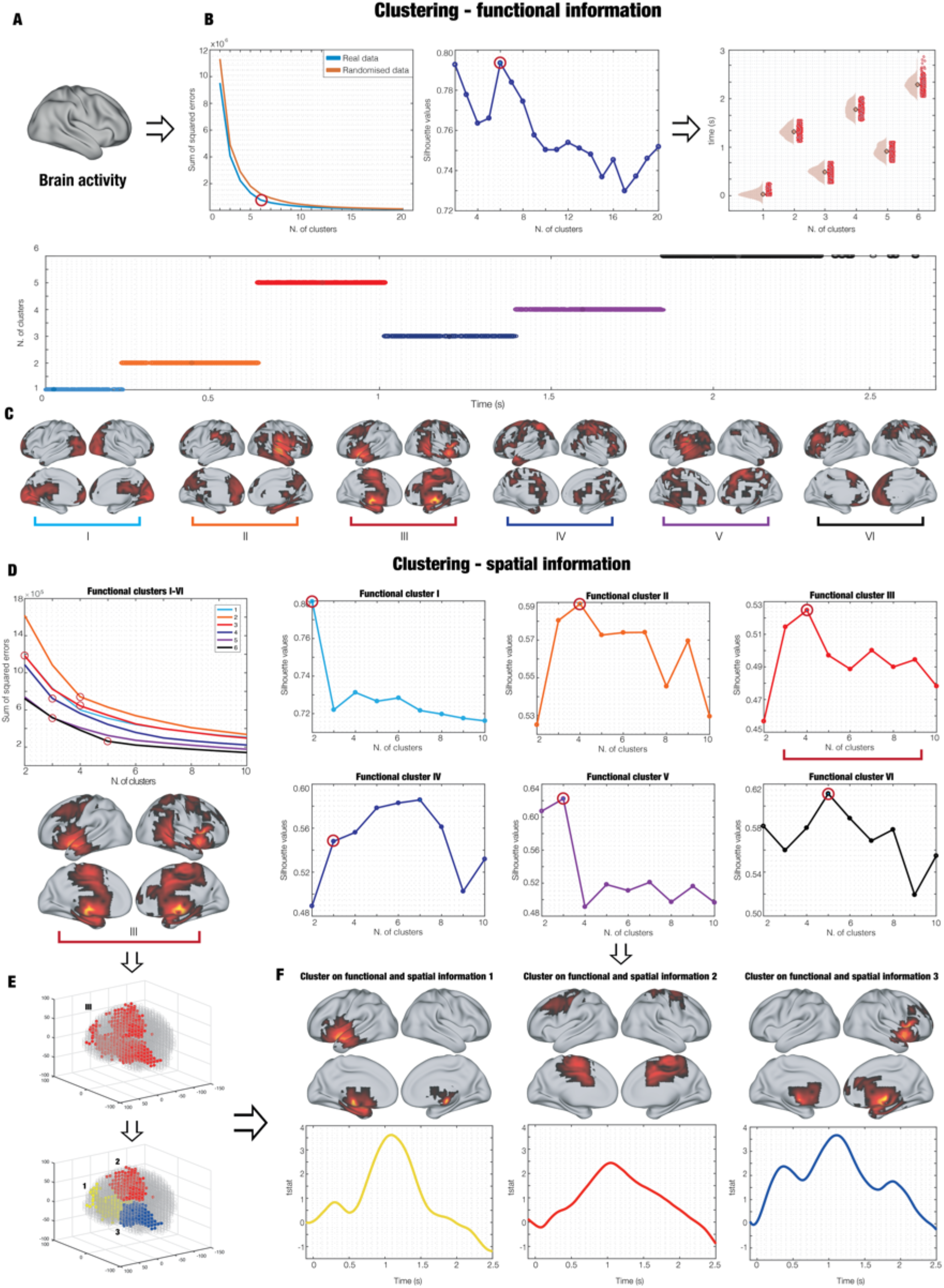
Description of the k-means functional clustering. (**A**) The brain activity is recorded during the recognition of temporal patterns. Such activity, reconstructed in 3559 brain sources, is the input for the k-means functional clustering to define a discrete functionally based parcellation. (**B**) A functional k-means clustering is performed. Such procedure consists of computing a series of *k* k-means clustering solutions (e.g. from *k* = 2 to *k* = 20) on the functional profile of the brain sources timeseries. In our study, we proposed two simple functional features: the time-index of the peak activity or the actual peak activity value of each brain source timeseries. The example reported in the figure shows clustering on time-indices of peak activity. The first plot shows the heuristic named elbow rule which helps to define the best *k* solution by plotting the sequential sum of squared errors (SSE) of the different cluster solutions (with *k* = 2:20). Here, it is visible how the SSE reduces its change rate around *k* = 6 (as indicated by the circle). Notably, when computing k-means clustering on randomized data, the SSE is higher for randomized vs real data, especially around *k* = 6, suggesting that the real data should be indeed clustered in six different clusters. As an alternative, the subsequent plot shows the Silhouette value for each *k*, representing how well each element (brain source time-index) is representative of the cluster to which it belongs. Ideally, those two heuristics should be considered together to define the best *k*. The plot on the right shows the brain source peak value indices in a violin-scatter fashion, while the plot below provides the same information with time on the x-axis and different colors for the six identified clusters to increase readability. (**C**) Brain representation of functional k-means clustering results (functional brain parcels). (**D**) A spatial k-means clustering is performed on each of the functional brain parcels to define a final parcellation considering both brain functional and spatial information. This procedure uses k-means clustering on the spatial coordinates of each of the brain sources belonging to each functional parcel (as shown especially by the plot of the elbow rule for all the six functional parcels). Then, to provide a specific example, the figure focuses on the third functional parcel (indicated by the red brace), whose plots for Silhouette heuristics are reported. (**E**) Graphical depiction of spatial parcels computation (bottom plot) for the third functional parcel (top plot). (**F**) Example of few final ‘k-means functional’ parcels and corresponding timeseries, obtained by averaging the timeseries of each brain source belonging to the parcel.

**Fig. S4.**
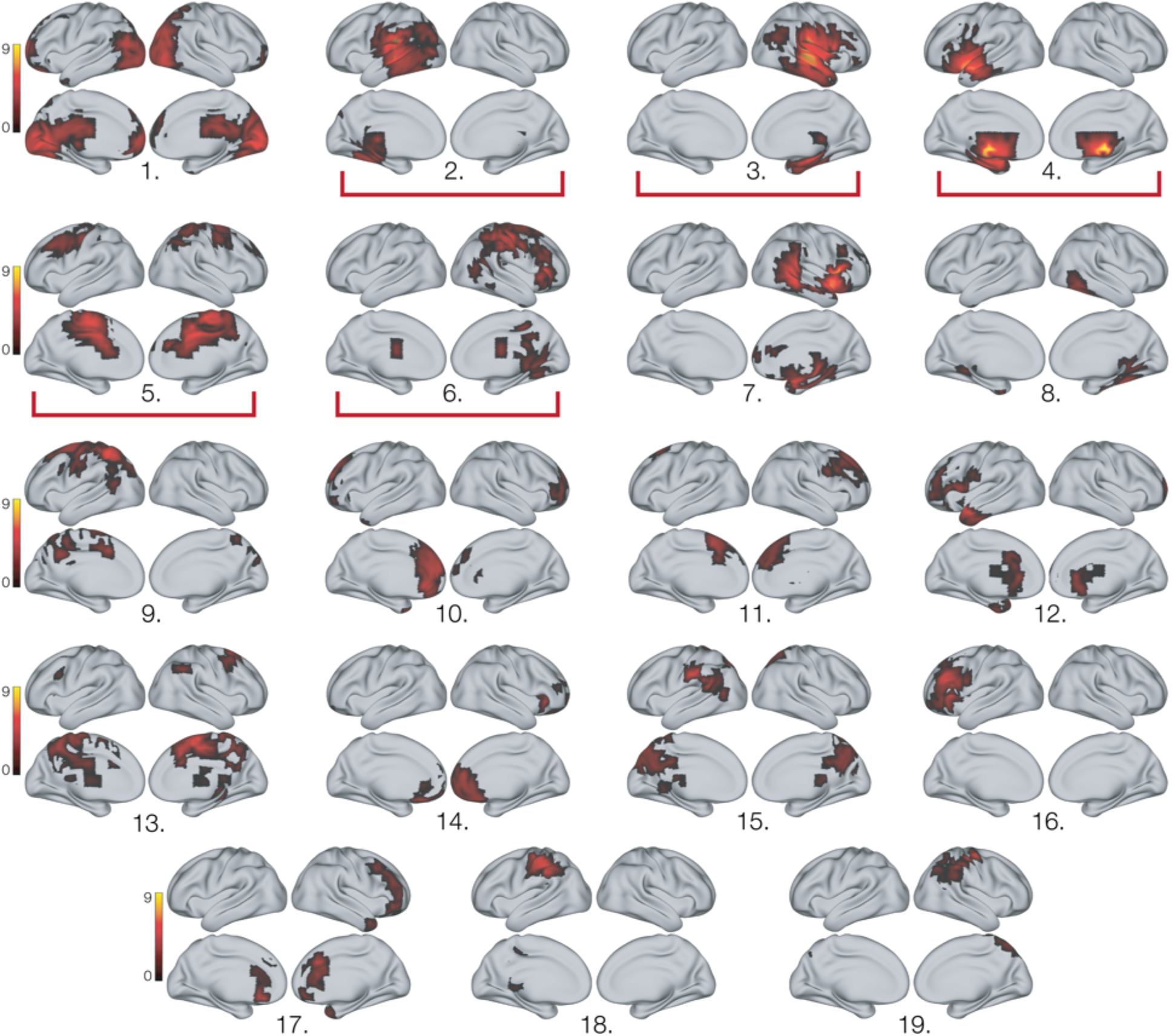
Functional parcellation for 0.1-1 Hz frequency band. Full parcellation returned by the k-means functional clustering computed on the indices of the brain activity peaks of all 3559 brain reconstructed sources. This parcellation was computed for the brain activity underlying recognition of memorized temporal patterns. The red brackets show the parcels that are reported in Fig. 2.

**Fig. S5.**
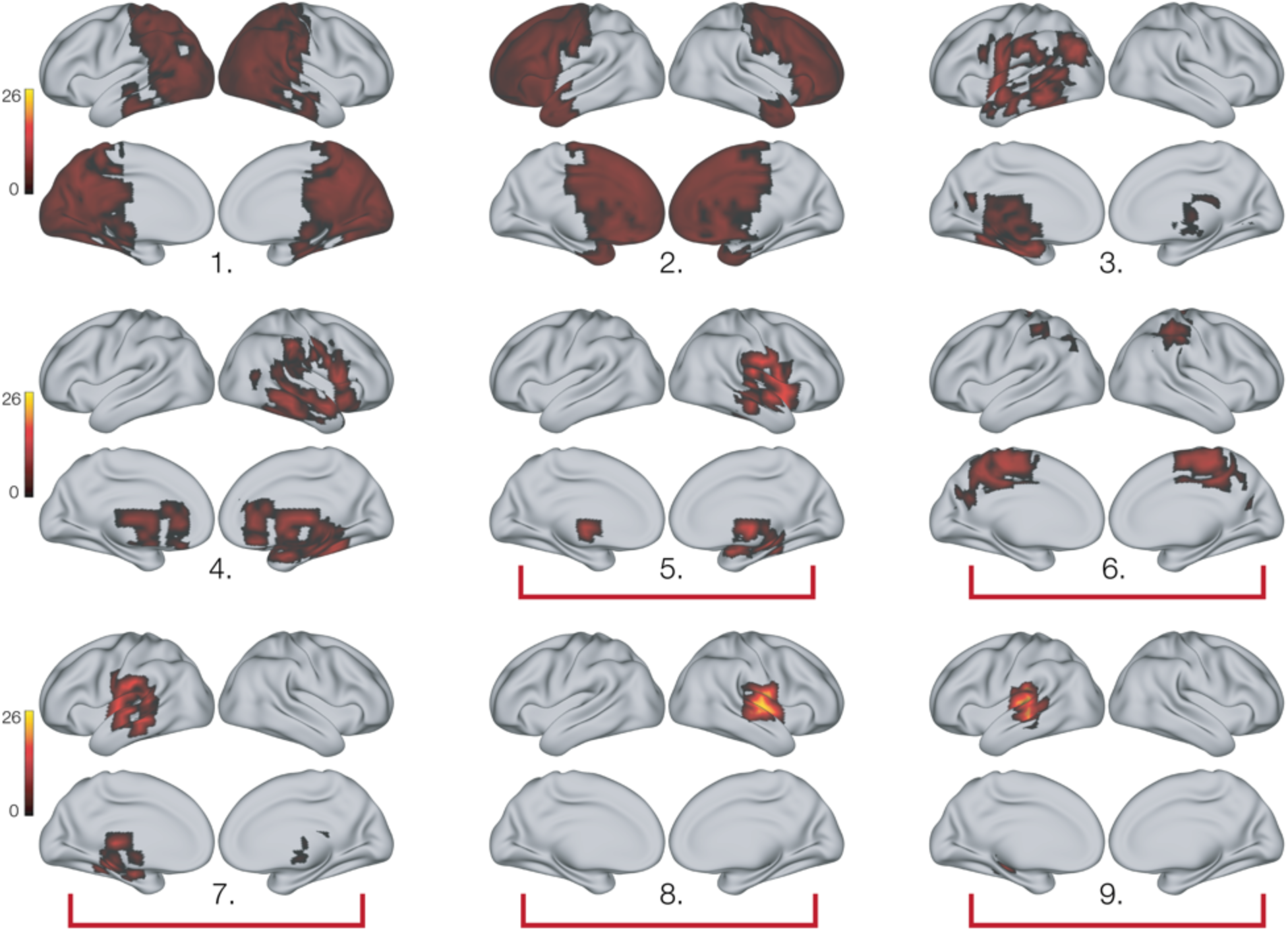
Functional parcellation for 2-8 Hz frequency band. Full parcellation returned by the k-means functional clustering computed on the brain activity peak values of the timeseries of all 3559 brain reconstructed sources. This parcellation was computed for the brain activity underlying recognition of memorized temporal patterns. The red brackets show the parcels that are reported in Fig. 2.

**Fig. S6.**
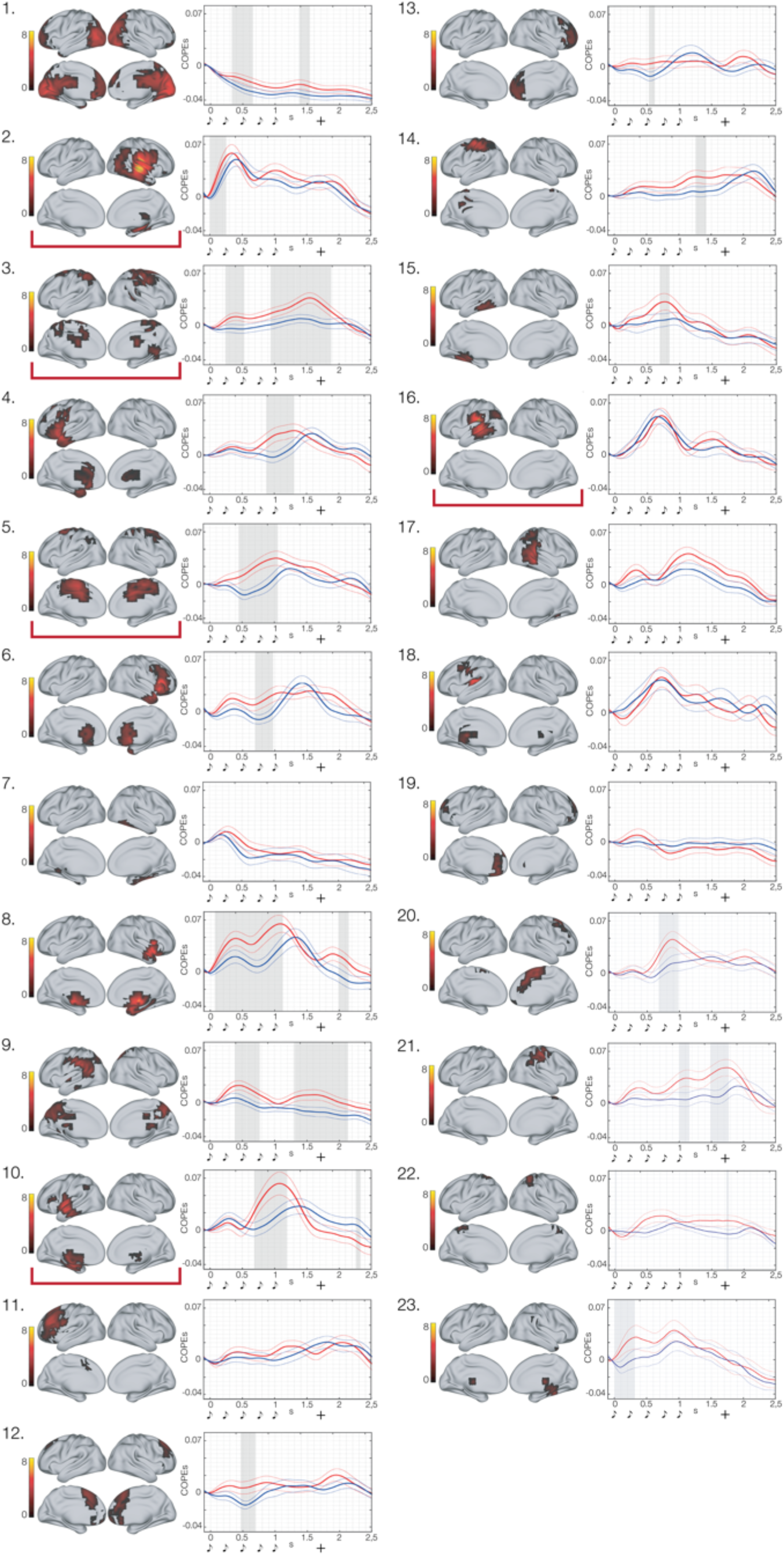
Full-parcellation contrasts between ‘memorized’ vs ‘novel’ temporal patterns in 0.1-1 Hz. Full parcellation and corresponding timeseries returned by the k-means functional clustering computed on the indices of the brain activity peaks of all 3559 brain reconstructed sources. In this case, the parcellation was computed for the averaged brain activity underlying recognition of ‘memorized’ and ‘novel’ temporal patterns. The brain parcels are numbered progressively with decreasing size (i.e. number of brain sources belonging to each parcel). The graphical depiction of musical tones indicates the onset of the objects forming the temporal pattern, while the ‘+’ shows the man reaction time of participants’ response. Grey areas illustrate the significantly different time-windows between M and N. In the waveform plots, the solid line corresponds to the mean brain activity, while the dash line to the correspondent standard errors. The red brackets show the parcels that are reported in Fig. 2.

**Fig. S7.**
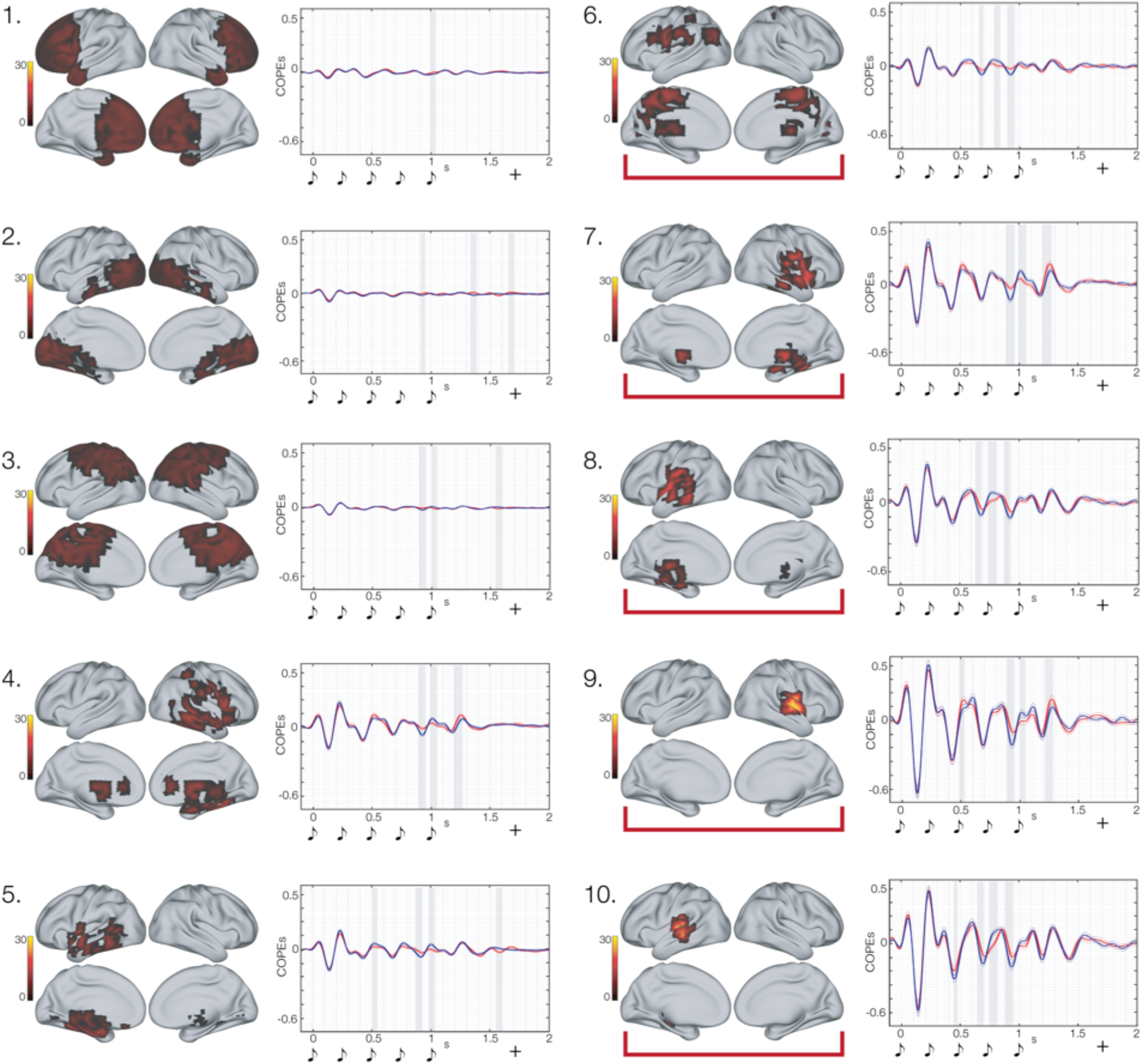
Full-parcellation contrasts between ‘novel’ vs ‘memorized’ temporal patterns in 2-8 Hz. Full parcellation and corresponding timeseries returned by the k-means functional clustering computed on the brain activity peak values of the timeseries of all 3559 brain reconstructed sources. In this case, the parcellation was computed for the averaged brain activity underlying recognition of ‘memorized’ and ‘novel’ temporal patterns. The brain parcels are numbered progressively with decreasing size (i.e. number of brain sources belonging to each parcel). The graphical depiction of musical tones indicates the onset of the objects forming the temporal pattern, while the ‘+’ shows the man reaction time of participants’ response. Grey areas illustrate the significantly different time-windows between N and M. In the waveform plots, the solid line corresponds to the mean brain activity, while the dash line to the correspondent standard errors. The red brackets show the parcels that are reported in Fig. 2.

**Fig. S8.**
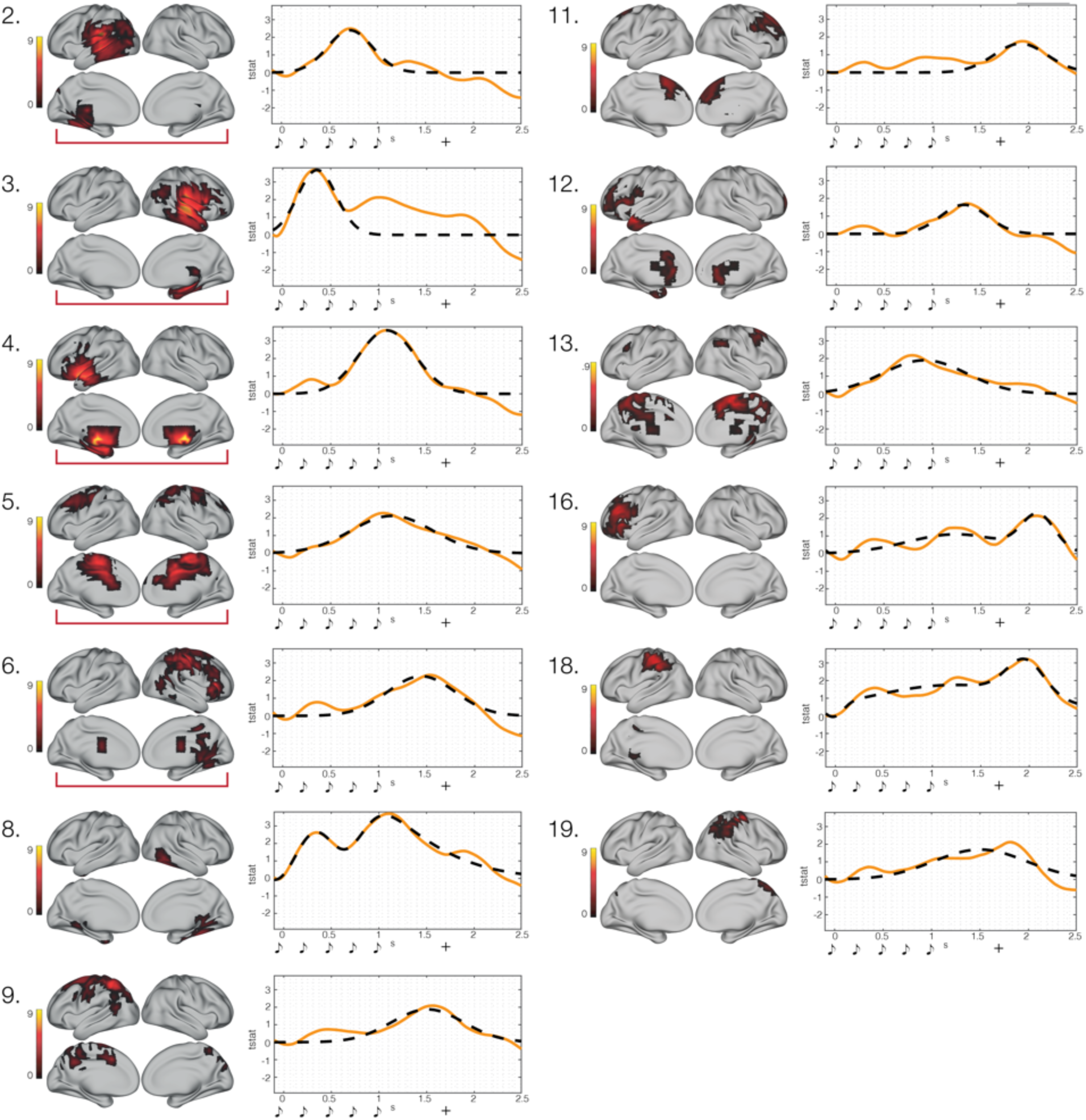
Full-parcellation fitting for ‘memorized’ temporal patterns in 0.1-1 Hz. All parcels whose timeseries were describable by a Gaussian function are reported in this figure. In a few cases, it was not possible to fit the equations since the timeseries showed a very small and scattered activity over time. This happened when those brain areas were not involved in the experimental task. For instance, this was the case of a large occipital parcel that, as conceivable, did not play any role in recognition of auditory sequences. The depicted parcels and corresponding timeseries were returned by the k-means functional clustering computed on the indices of the brain activity peaks of all 3559 brain reconstructed sources. This parcellation was computed for the brain activity underlying recognition of memorized temporal patterns (see Methods for details). The brain parcels are numbered progressively with decreasing size (i.e. number of brain sources belonging to each parcel). The graphical depiction of musical tones indicates the onset of the objects forming the temporal pattern, while the ‘+’ shows the man reaction time of participants’ response. In the waveform plots, the solid line corresponds to the actual brain activity, while the dash line to the predicted timeseries obtained by using non-linear least square fitting. The red brackets show the parcels that are reported in Fig. 2.

**Fig. S9.**
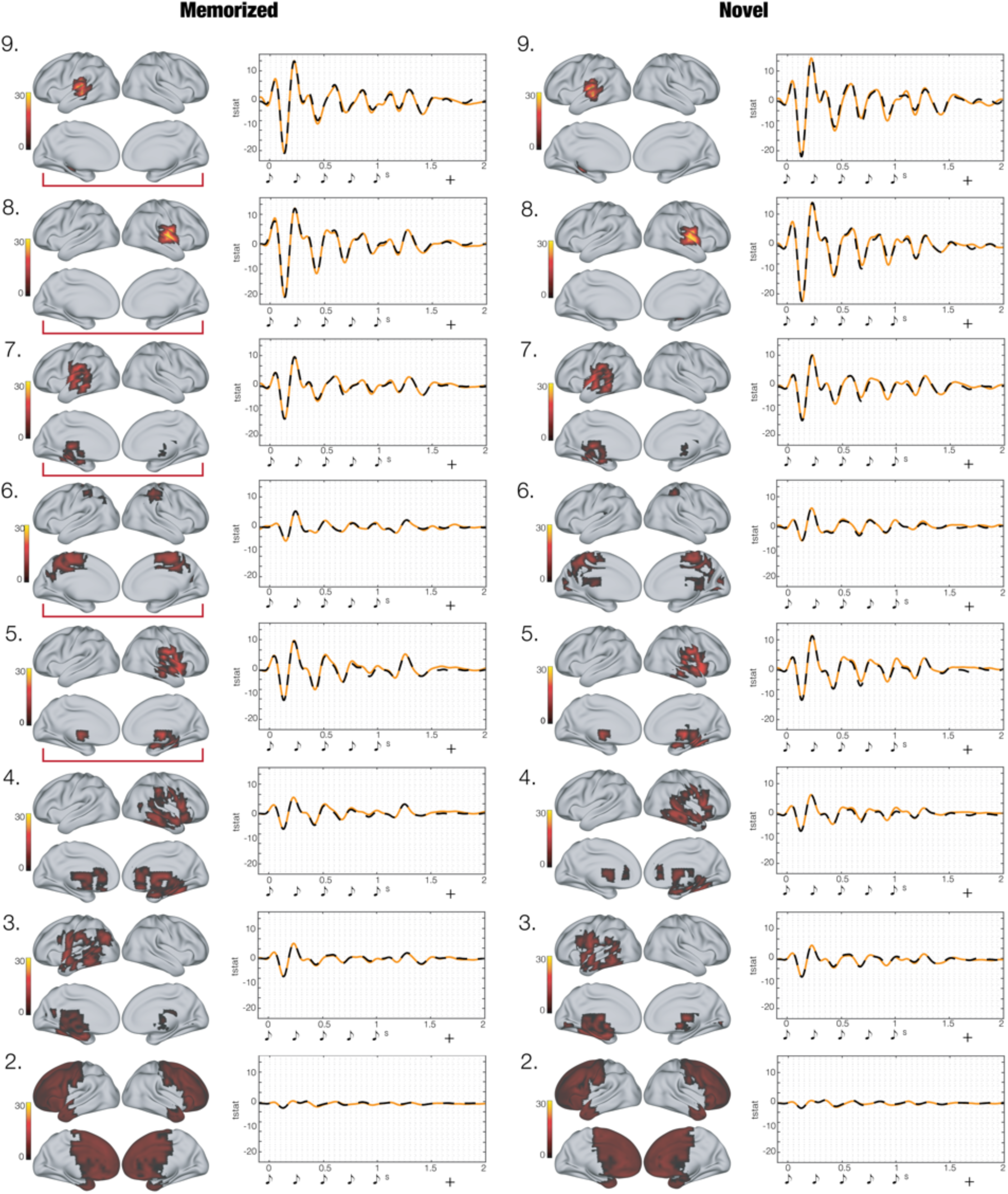
Full-parcellation fitting for ‘memorized’ and ‘novel’ temporal patterns in 2-8 Hz. All parcels whose timeseries were describable by a skewed Gaussian function multiplied by a sinusoidal function are reported in this figure. Only in one case which regarded a large occipital parcel, it was not possible to fit the equation since the timeseries showed a very small and scattered activity over time. This happened since, as conceivable, the occipital cortex did not play any role in the processing and recognition of auditory sequences. The depicted parcels and corresponding timeseries were returned by the k-means functional clustering computed on the brain activity peak values of the timeseries of all 3559 brain reconstructed sources. The two parcellations reported in the figure were computed for the brain activity underlying the recognition of either the ‘memorized’ (left column) or the ‘novel’ temporal patterns (right column) (see Methods for details). The brain parcels are numbered progressively with decreasing size (i.e. number of brain sources belonging to each parcel). The graphical depiction of musical tones indicates the onset of the objects forming the temporal pattern, while the ‘+’ shows the man reaction time of participants’ response. In the waveform plots, the solid line corresponds to the actual brain activity, while the dash line to the predicted timeseries obtained by using non-linear least square fitting. The red brackets show the parcels that are reported in Fig. 2.

## SUPPLEMENTARY TABLES

**Table S1. Brain activity underlying recognition of temporal patterns (single-object)**

Extensive brain sources activity underlying recognition of each object (musical tone) of the temporal patterns. Results are reported for recognition of ‘memorized’ (M) and ‘novel’ (N) sequences independently as well as for their contrasts. Brain areas refer to the automatic anatomic labelling (AAL) parcellation labels, while t indicates the t-value obtained by contrasting M vs N temporal sequences.

**Table S2. Functionally-based parcellation for recognition of ‘memorized’ patterns – 0.1-1 Hz**

Description of the brain sources belonging to each of the parcels returned by the k-means functional clustering. For each source, the table reports a descriptive label (referring to automatic anatomic labelling (AAL) parcellation), hemisphere, MNI coordinates, and maximum t-value registered in the source timeseries.

**Table S3. Functionally-based parcellation for recognition of ‘memorized’ and ‘novel’ patterns – 0.1-1 Hz**

Description of the brain sources belonging to each of the parcels returned by the k-means functional clustering. In this case, the clustering algorithm has been performed on the brain activity averaged over experimental conditions (‘memorized’ and ‘novel’). For each source, the table reports a descriptive label (referring to automatic anatomic labelling (AAL) parcellation), hemisphere, MNI coordinates, and maximum t-value registered in the source timeseries.

**Table S4. Functionally-based parcellation for recognition of ‘memorized’ patterns – 2-8 Hz**

Description of the brain sources belonging to each of the parcels returned by the k-means functional clustering performed for ‘memorized’ patterns. For each source, the table reports a descriptive label (referring to automatic anatomic labelling (AAL) parcellation), hemisphere, MNI coordinates, and maximum t-value registered in the source timeseries.

**Table S5. Functionally-based parcellation for recognition of ‘novel’ patterns – 2-8 Hz**

Description of the brain sources belonging to each of the parcels returned by the k-means functional clustering performed for ‘novel’ patterns. For each source, the table reports a descriptive label (referring to automatic anatomic labelling (AAL) parcellation), hemisphere, MNI coordinates, and maximum t-value registered in the source timeseries.

**Table S6. Functionally-based parcellation for recognition of ‘memorized’ and ‘novel’ patterns – 2-8 Hz**

Description of the brain sources belonging to each of the parcels returned by the k-means functional clustering. In this case, the clustering algorithm has been performed on the brain activity averaged over experimental conditions (‘memorized’ and ‘novel’). For each source, the table reports a descriptive label (referring to automatic anatomic labelling (AAL) parcellation), hemisphere, MNI coordinates, and maximum t-value registered in the source timeseries.

**Table S7. Brain activity underlying recognition of temporal patterns (k-means functional clustering)**

Contrasts between brain activity underlying recognition of ‘memorized’ vs ‘novel’ temporal patterns. Here, the contrasts have been performed on the timeseries of the parcels returned by the k-means functional clustering computed on the brain activity averaged over experimental conditions. The table provides results for both frequencies (0.1-1. Hz and 2-8 Hz). Further, for each parcel, it reports size (*k*), *p-value* corrected by Monte-Carlo simulations, temporal extent, and averaged *t-value* of the significant clusters.

**Table S8. Fitted coefficients over all parcels’ timeseries (non-linear least square)**

R^2^ and coefficients derived from non-linear least square fitting of the equations (5), (6) and (7) reported in the Methods section on the brain activity underlying temporal pattern recognition. In a few cases, it was not possible to fit the equation since the timeseries showed a very small and scattered activity over time. This happened when those brain areas were not involved in the experimental task. For instance, this was the case of a large occipital parcel that, as conceivable, did not play any role in recognition of auditory sequences. The reported parcels were returned by the k-means functional clustering computed either on the indices or on the actual brain activity maximum values of all 3559 brain reconstructed sources. This parcellation was computed for the brain activity underlying recognition of ‘memorized’ temporal patterns for 0.1-1 Hz and ‘novel’ temporal patterns for 2-8 Hz only (see Methods for details).

